# A paradoxical population structure of *var* DBLα types in Africa

**DOI:** 10.1101/2023.11.05.565723

**Authors:** Mun Hua Tan, Kathryn E. Tiedje, Qian Feng, Qi Zhan, Mercedes Pascual, Heejung Shim, Yao-ban Chan, Karen P. Day

## Abstract

The *var* multigene family encodes the *P. falciparum* erythrocyte membrane protein 1 (PfEMP1), which is important in host-parasite interaction as a virulence factor and major surface antigen of the blood stages of the parasite, responsible for maintaining chronic infection. Whilst important in the biology of *P. falciparum*, these genes (50 to 60 genes per parasite genome) are routinely excluded from whole genome analyses due to their hyper-diversity, achieved primarily through recombination. The PfEMP1 head structure almost always consists of a DBLα-CIDR tandem. Categorised into different groups (upsA, upsB, upsC), different head structures have been associated with different ligand-binding affinities and disease severities. We study how conserved individual DBLα types are at the country, regional, and local scales in Sub-Saharan Africa. Using publicly-available sequence datasets and a novel ups classification algorithm, *cUps*, we performed an *in silico* exploration of DBLα conservation through time and space in Africa. In all three ups groups, the population structure of DBLα types in Africa consists of variants occurring at rare, low, moderate, and high frequencies. Non-rare variants were found to be temporally stable in a local area in endemic Ghana. When inspected across different geographical scales, we report different levels of conservation; while some DBLα types were consistently found in high frequencies in multiple African countries, others were conserved only locally, signifying local preservation of specific types. Underlying this population pattern is the composition of DBLα types within each isolate DBLα repertoire, revealed to also consist of a mix of types found at rare, low, moderate, and high frequencies in the population. We further discuss the adaptive forces and balancing selection, including host genetic factors, potentially shaping the evolution and diversity of DBLα types in Africa.

## 1. INTRODUCTION

Malaria parasites in endemic areas with high transmission undergo frequent outcrossing in the vector (1,2). The diversification of parasites through meiotic, mitotic, and ectopic recombination results in very high levels of genetic diversity in the *Plasmodium falciparum* parasite population (3–7), particularly of the major variant surface antigens such as the *var* multigene family that encode the *Plasmodium falciparum* erythrocyte membrane protein 1 (PfEMP1). PfEMP1 proteins are expressed on the surface of infected erythrocytes and can bind to host receptors to mediate cytoadhesion and sequestration of infected cells (8–10). Through clonal antigenic variation of *var* genes, whereby different genes are expressed sequentially and exclusively during the blood stage, parasites are also able to effectively evade immune detection, promoting chronic infection within a host (11,12). This high diversity has served as the basis for *var* surveillance (13,14), population genetics (15–17), and estimation of infection complexity (15). However, less work has been done to characterise this diversity and the population structure of antigenic factors of and within these genes.

There are approximately 40 to 60 different *var* genes found across all 14 chromosomes of a *P. falciparum* genome (18). Based on their upstream promoter sequences, *var* genes can be divided into groups of A, B, C, and E, with a minority of genes grouped in two intermediate groups of B/A or B/C (19). The three major ‘ups’ groups of upsA, upsB, and upsC are associated with different chromosomal locations, transcriptional directions, and sequences (18–21). Genes in the upsA and upsB groups are generally located at the subtelomeric regions whereas genes in the upsC group are found in the central regions of specific chromosomes. UpsA genes have been shown to be transcribed towards centromeres and conversely, upsB and upsC genes are often found to be transcribed towards telomeres.

The extracellular N-terminal PfEMP1 head structure almost always consists of a Duffy-binding-like alpha domain (DBLα) and a cysteine-rich interdomain region (CIDR) (i.e., a DBLα-CIDR tandem). This head structure exists in different configurations of these DBLα and CIDR domain subclasses (e.g., DBLα0-2 with CIDRα1-6, CIDRβ1-7, CIDRδ1-12, CIDRγ1,2) and can influence ligand binding and disease pathogenicity. The prevailing understanding is that *var* genes and DBLα variants (i.e., DBLα types) in the upsA group are generally more conserved compared to those in upsB or upsC groups (i.e., non-upsA) (15,16,19,22). This has largely been attributed to the association of upsA genes to severe malaria, including cerebral malaria, due to their host receptor binding phenotypes (23). Expression of upsA *var* genes encoding the DBLα+CIDRα1 head structure mediate endothelial protein C receptor (EPCR)-binding and/or intercellular adhesion molecule-1 (ICAM-1)-binding has been associated with severe malaria and/or cerebral malaria (24–26). In addition, PfEMP1 in the upsA group containing the DBLα+CIDRβ/δ/γ head structure has been associated with rosetting with uninfected erythrocytes (24). On the other hand, expression of upsB and upsC *var* genes (e.g., DBLα+CIDRα2-6) have been associated with uncomplicated malaria, commonly mediated by adhesion to the cluster differentiation 36 (CD36) receptor (27,28), and may be more active in establishing chronic infections (29).

With the exception of one specific *var* gene involved in pregnancy-associated malaria (i.e., *var2csa*), all other *var* genes encode for a DBLα domain, which is one of the most diverse domains and has been shown to be immunogenic to variant-specific epitopes, recognised serologically in an age-dependent manner (30). Multiple studies have noted that there exists a minority of DBLα types that are highly conserved over various spatial scales (16,31–35). These studies typically focussed on DBLα types or *var* genes found within the highest percentiles (e.g., (35)), at very high frequencies (e.g., (16,31)), or those found globally conserved and prevalent (e.g., (34,35)). Understandably, looking for the most common DBLα or *var* is instinctive in the search for the elusive vaccine candidate targeting the most important group of variant surface antigens of the parasite. However, *var* gene sharing among isolates, particularly those living in high transmission, has been shown to be minimal (33). Given that multiple variants of *var* genes with the same binding phenotype exist, this pre-occupation over the most globally-common types or genes risks overlooking the genetic patterns underlying a local population.

In the same way that ‘severe malaria’ must defined by different malaria epidemiologies, patterns of conservation must also be interpreted within the context of an area’s local epidemiology. In low-transmission areas in South America and Asia, conservation found across countries and continents likely relates to small population sizes due to founder effects, in which *var* genes have not yet diversified (36–38). In moderate transmission, profiles may exhibit bias toward more moderate frequency classes, in conjunction with greater diversity within the area. Conservation in high-transmission areas would be most interesting, as these areas possess the epidemiological and genetic characteristics to generate vast diversity. In such a system of high prevalence/incidence, high outcrossing rates, and high genetic diversity of parasites, finding conservation will provide insights into factors constraining and shaping diversity in a highly dynamic system.

Equipped with publicly-available DBLα data from several populations in Africa and a novel ups classification algorithm (***cUps***) introduced in this paper, this study sought to identify DBLα type conservation beyond those reported in the highest frequencies within and among populations globally. By categorising DBLα types into broad frequency classes of rare to highly-frequent types, we identified two kinds of conservation within all ups groups in our study area in Bongo, Ghana: (1) Conservation of specific types across isolates in a time point and through time (i.e., survey), (2) Conservation of type frequencies (i.e., types were found at relatively stable frequencies through time). We show that these patterns are maintained through the composition of DBLα types within each isolate repertoire, revealed to consist of a mix of types found at rare, low, moderate, and high frequencies in the population. In a spatial analysis, in addition to identifying DBLα types conserved at the continent level, we noted that there are DBLα types conserved at the local and/or regional levels but not necessarily prevalent across wider geographical scales, prompting a discussion on the adaptive forces potentially driving balancing selection and shaping the population structure of DBLα types in Africa.

## 2. RESULTS

### 2.1 Description of time-series cross-sectional surveys in Bongo, Ghana

This study analysed publicly-available DBLα tag sequence data from an interrupted time-series study design (i.e., Malaria Reservoir Study (MRS)) (Figure I in Data S1) (13,15,39). This MRS dataset consists of sampling at seven time points from 2012 to 2017 at the end of wet or dry seasons. Each time point represented an age-stratified cross-sectional survey of approximately 2,000 asymptomatic participants per survey (ages from 1 to 97 years old) from two proximal catchment areas (Vea/Gowrie and Soe, with a sampling area ∼60 km^2^) in Bongo District in Northern Ghana. Surveyed participants (i.e., isolates) represented approximately 15% of the total population that reside in the two catchment areas in Bongo District at a time (Table I in Data S1). This area is characterised by high, seasonal malaria transmission and has undergone several types of interventions, including long-lasting insecticide-treated nets (LLINs) and indoor residual spraying (IRS) that reduced transmission (13,15), and seasonal malaria chemoprevention (SMC) that reduced the burden of infection in children younger than 5 years old (15). Clustering of DBLα tag sequences from seven surveys (S1 to S7) and further post-processing of the dataset (see Methods) resulted in 62,168 representative DBLα types found in 3,166 isolates for this study of DBLα conservation in Bongo (Table I in Data S1).

In a high-transmission setting, the asymptomatic “population” typically consists of “isolates” infected by one or more unique parasite “genomes”. This complexity of infections is indicated by multiplicity of infection (MOI), where an isolate with MOI = 1 would represent a single unique parasite genome. Hence, at MOI = 1, an isolate’s DBLα repertoire is synonymous to a parasite’s DBLα repertoire (i.e., the DBLα repertoire in a single parasite genome). Conversely, at MOI > 1, an isolate’s DBLα repertoire would encompass > 1 parasites’ DBLα repertoire.

### 2.2 A novel ups classification algorithm based on *var* DBLα tags

This study introduces ***cUps***, a novel algorithm for classifying DBLα types into the different groups of upsA, upsB, and upsC (Data S2, Figure 1A). At the isolate level, the average isolate repertoire consists of 20.9%, 48.6%, and 30.5% of upsA, upsB, and upsC DBLα types, respectively (Figure II in Data S1). These proportions differ from those reported in (22) that estimated higher proportions of upsB and lower proportions of upsC in isolate repertoires, based on the average of seven genomes. The algorithm shows a tendency to classify more upsB types as upsC types. This is in line with validation results on the algorithm’s specificity and sensitivity presented Data S2. A reduced analysis that involved DBLα types with higher confidence in classification (threshold = 8, see Data S2) yielded similar trends and patterns of observation. Genetic similarity by pairwise type sharing (PTS) remains extremely low for all ups groups (median PTS: 4.55% (upsA), 1.00% (upsB), and 2.15% (upsC)) (Figure 1B). The 62,168 representative DBLα types from the seven combined MRS surveys were classified into upsA (5.4%), upsB (56.6%), and upsC (37.9%) groups (Table I in Data S1). This points to a highest DBLα richness for the upsB group (35,215 types), followed by the upsC group (23,583 types) and upsA group (3,370 types) in combined surveys, and this hierarchy of richness is similarly observed for individual surveys (Figure 1A). The differences in proportions of ups groups at the isolate versus population levels is attributed to the negative relationship between PTS and richness, as a higher level of upsA type sharing will result in lower proportion of unique representative DBLα types in the population.

**Figure 1.**
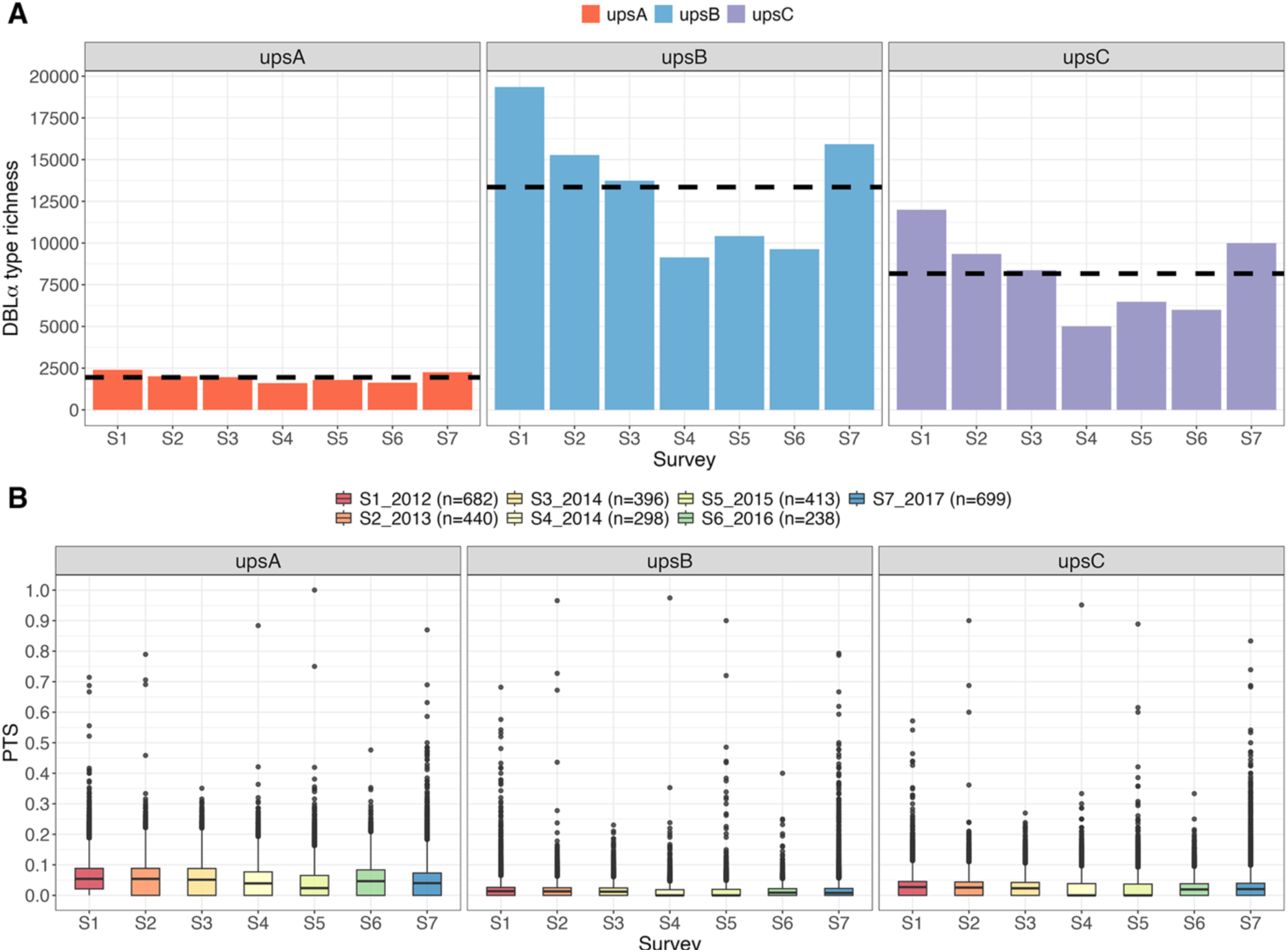
**Classification of DBLα types into ups groups (upsA, upsB, upsC) [Malaria Reservoir Study (MRS)].** Figure shows (A) DBLα richness (horizontal dashed lines show mean richness per ups group) and (B) genetic similarity (i.e., overlaps in isolate repertoire by pairwise type sharing (PTS)), in each of the seven MRS surveys in Bongo, Ghana.

### 2.3 *Var* DBLα types are conserved in the local Bongo population and through time

The frequency of a DBLα type is defined as the proportion of the population a DBLα type is found in and is calculated in the context of individual surveys (i.e., “survey-specific frequency”), or of the averaged frequencies of the seven surveys (i.e., “survey-averaged frequency”, see Methods for details on frequency calculation and normalisation). To explore the conservation of DBLα types, we further categorised these frequencies into four classes of ***low*** (0%, 1%), ***moderate*** [1%, 5%), ***high*** [5%, 10%), and ***very high*** [10%, 100%] frequencies. While it is well reported that DBLα types in the upsA group are generally more conserved relative to types in the upsB and upsC groups, this study identified conservation of DBLα types in all three ups groups. We describe patterns of DBLα conservation observed in Bongo at the population level in the following subsections.

#### 2.3.1. Conservation of var DBLα types in a surveyed time point

DBLα types are considered conserved in a surveyed time point when found in multiple isolates sampled in a same survey (i.e., moderate-to-high survey-specific frequencies). Overall, distributions of survey-averaged and survey-specific frequencies were found to be strongly and positively skewed, with most DBLα types occurring at low frequencies in <1% of isolates. For all three ups groups, when categorised into frequency classes, the second largest subsets of DBLα types are shown to occur at moderate frequencies between 1% to 5% in each survey, followed by smaller subsets of DBLα types found at higher frequencies exceeding 5% and/or 10% frequencies (Figure 2A). The most frequent DBLα types in the upsA, upsB, and upsC groups were detected at survey-specific frequencies of 61.1%, 42.9%, and 62.0%, respectively. In the different surveys, this study identified hundreds to thousands of moderate-to-highly conserved DBLα types (i.e., ≥1% survey-specific frequencies) in all three ups groups (upsA: 525 to 790 types per survey, upsB: 539 to 1,917 types per survey, upsC: 365 to 1,121 types per survey). This translates into different proportions of DBLα types in each ups group, owing to the higher DBLα type richness of upsB and upsC groups (upsA: 29.2% to 43.9% per survey, upsB: 5.2% to 15.5% per survey, upsC: 5.6% to 14.3% per survey) (Table II in Data S1).

**Figure 2.**
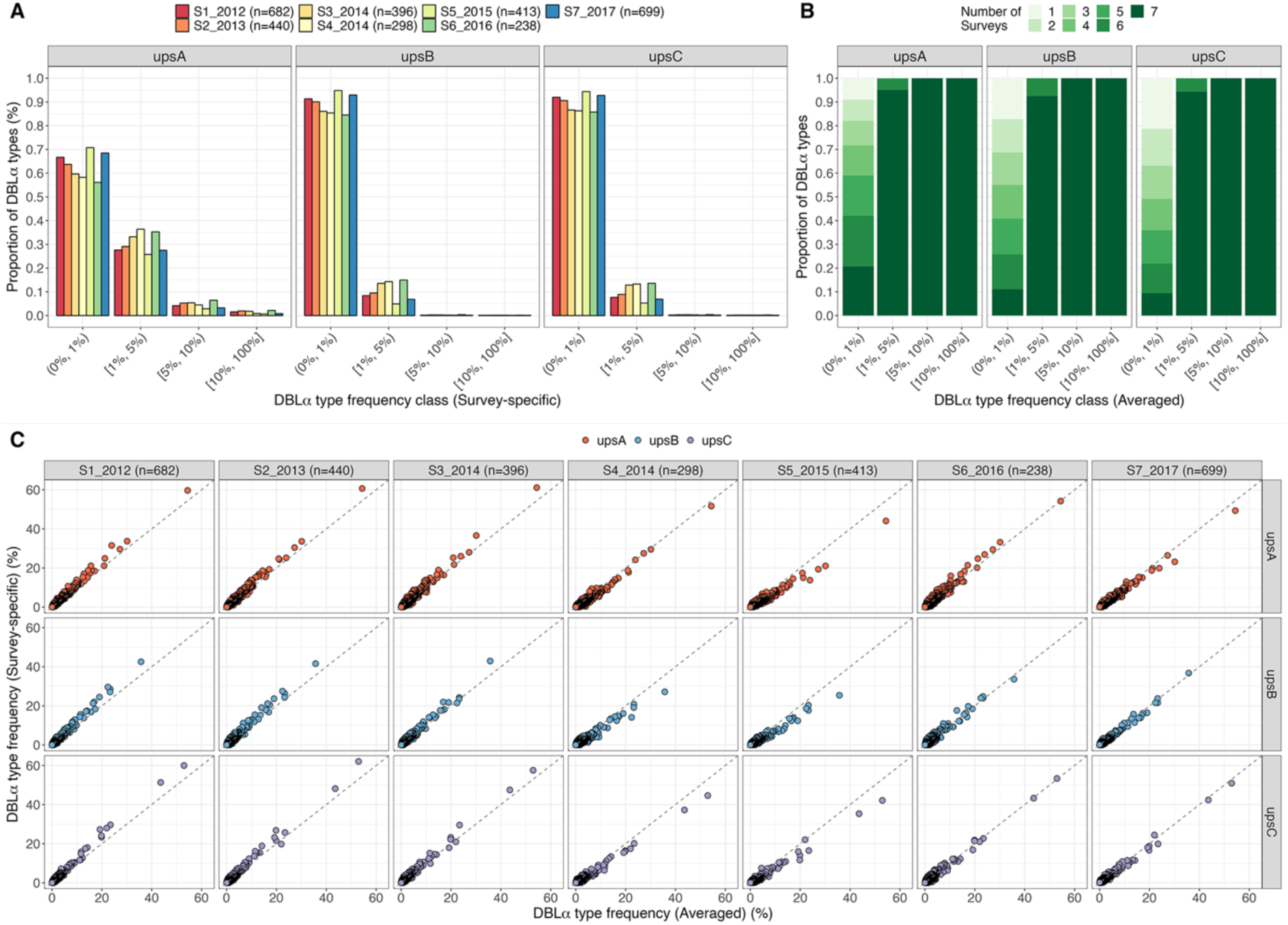
**Conservation of DBLα types and frequencies in a local population and through time [Malaria Reservoir Study (MRS)].** (A) Distribution of survey-specific frequencies of DBLα types, binned into categorical frequency classes. (B) Number of surveys DBLα types were observed in, showing that DBLα types found at ≥1% survey-averaged frequencies were seen to also persist through most surveys. This is also true for DBLα types found at ≥ 1% survey-specific frequencies (Figure IV in Data S1). (C) Survey-specific frequencies (y-axis) of DBLα types are plot against survey-averaged frequencies (x-axis), showing positive correlation between both frequencies. Points represent individual DBLα types, coloured by ups groups.

#### 2.3.2. Conservation of var DBLα type frequencies across multiple surveys and through time

Frequencies of DBLα types are considered conserved if a same DBLα type occurs at stable or similar frequencies in multiple surveys. In all ups groups, a strong positive correlation between survey-specific frequencies and survey-averaged frequencies of DBLα types is shown, indicating that DBLα types occurring at high survey-averaged frequencies were also generally found at high frequencies in specific surveys and likely in multiple surveys (Figure 2C, Figure III in Data S1). Likewise, most DBLα types found at moderate frequencies also showed consistent maintenance of frequencies across multiple surveys. Furthermore, DBLα types are considered conserved through time when found in multiple surveys. This study shows a correlation between DBLα type frequencies (per-survey and survey-averaged) and the number of surveys the types are found in, with most of the relatively conserved DBLα types in all three ups groups also persisting through time (Figure 2B, Figure III and IV in Data S1). All DBLα types occurring at high or very high survey-averaged and/or survey-specific frequencies were seen in all seven surveys. The majority of those with moderate survey-averaged or survey-specific frequencies were found in all seven surveys, with a smaller subset found in four to six surveys. On the other hand, the remainder of DBLα types found at low frequencies (< 1%) can be found in the range of one to seven surveys. This was observed in all three ups groups.

### 2.4 Isolate repertoires consist of a mix of conserved and rare *var* DBLα types

While isolate DBLα repertoires in high-transmission populations have been reported to be unrelated and largely non-overlapping (15,17), there has not been a detailed exploration of the composition of DBLα types and their respective frequency classes at the isolate repertoire level (i.e., ‘per-isolate frequency profiles’). This study showed that all isolate repertoires consist of DBLα types from all three ups groups. More interestingly, these per-isolate frequency profiles show consistent proportions of rare, moderately-frequent, and highly-frequent DBLα types in every isolate, based on survey-specific frequencies (Figure 3, Figure V in Data S1). In all ups groups, these frequency profiles were consistent across isolates within the same survey, regardless of isolates’ infection complexities (i.e., MOI). Importantly, the observation of these frequency patterns in MOI = 1 isolates (i.e., isolate non-upsA repertoire size of approximately ≤ 45 in Figure 3) indicates that these per-isolate frequency profiles are a consequence of similar repertoire composition within actual parasite genomes.

**Figure 3.**
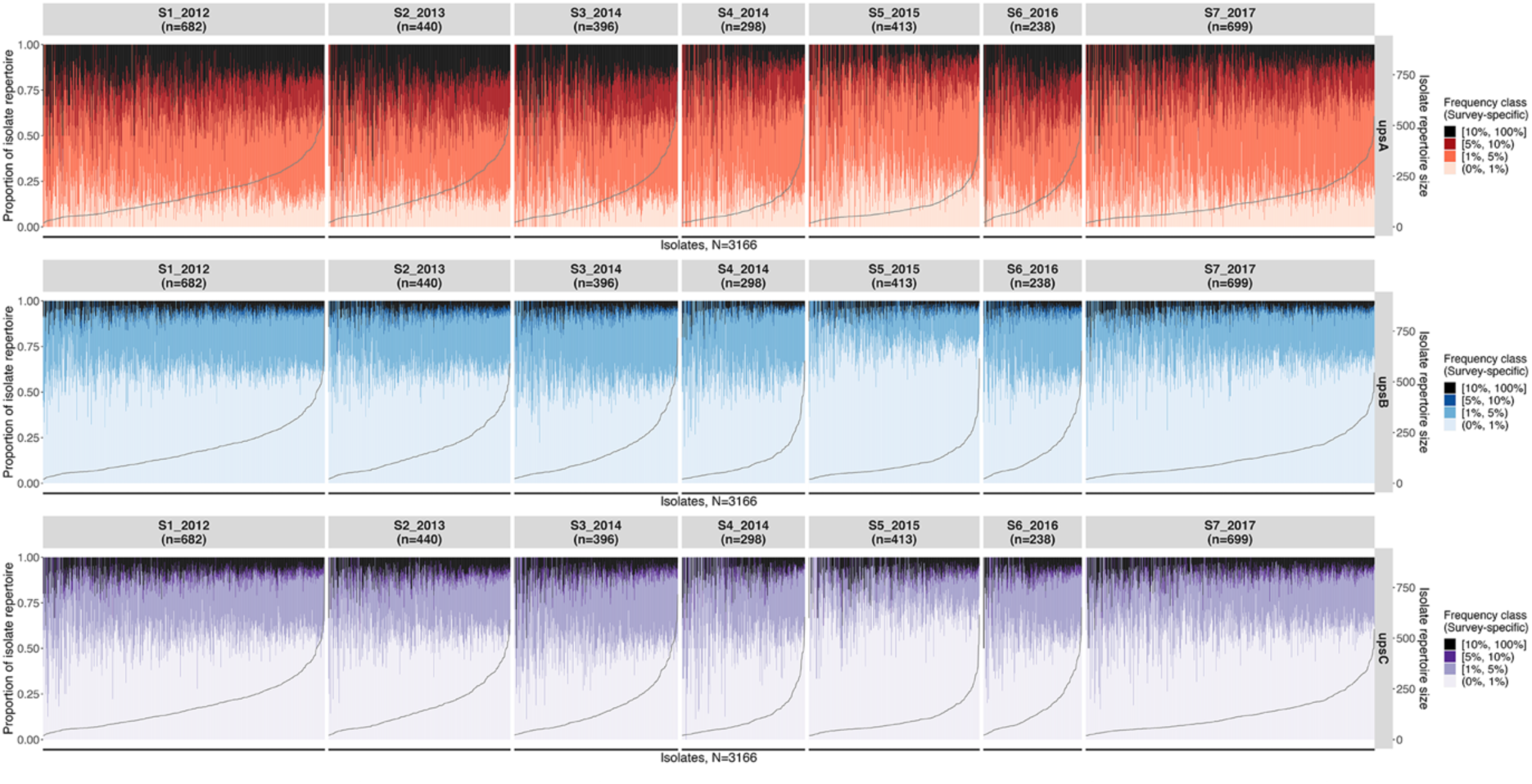
**Per-isolate frequency profiles show the composition of survey-specific frequency classes in every isolate repertoire [Malaria Reservoir Study (MRS)].** Vertical bars represent individual isolates, showing the composition of frequency classes in each isolate (left y-axis) by ups group (horizontal panels). Isolates are sorted by isolate repertoire size in increasing order, with isolate repertoire sizes indicated by the grey line (right y-axis).

Of the three groups, isolates’ upsA frequency profiles exhibit largest proportions of DBLα types found at moderate survey-specific frequencies and smaller proportions of DBLα types in the lowest survey-specific frequency class. In contrast, isolates’ upsB and upsC frequency profiles both consist of largest proportions of DBLα types in the lowest frequency class (i.e., (0%, 1%)), followed by those in the moderate frequency class. The introduction of interventions did not perturb these frequency profiles, which maintained the composition of different frequency classes, albeit in different proportions. In surveys of the population affected by interventions (e.g., S4 and S5 during and after indoor residual spraying), frequency profiles trended toward a generally larger proportion of low-frequency DBLα types and smaller proportions of higher frequency DBLα types within each isolate (Figure 3). We confirmed this observation of per-isolate frequency profiles using an independent DBLα sequence dataset (14,33,40), extracted from *var* genes of isolates sampled from Navrongo in Ghana, situated ∼30 km adjacent to Bongo district (Figure VI in Data S1).

As expected, genetic similarity between pairwise isolate repertoires (i.e., PTS values) increases when DBLα types of lower frequencies are excluded (Figure VII in Data S1). PTS values range from 0 to 1 representing unrelated to identical isolate repertoires, respectively. Interestingly, even when considering only DBLα types found at very high frequencies, median PTS values remained generally low (median PTS of 0.02, 0.05, 0.13, and 0.19 when considering DBLα types at >0%, ≥1%, ≥5%, ≥10% survey-averaged frequencies, respectively, across all surveys). This indicates that, while every isolate repertoire contains sets of types that are conserved in the population, identical sets of conserved DBLα types are rare. Shifts in PTS distributions were more substantial when exclusively evaluating DBLα types in the upsA or upsC groups relative to the upsB group, consistent with the lower DBLα richness in the two former groups (i.e., the less variants there are in the population, the higher the probability of overlaps).

### 2.5 Global and local preservation of *var* DBLα types in Africa

Further, a separate spatial study of DBLα conservation in multiple African countries (i.e., “locations”) representing West Africa (Ghana, Gabon), Central Africa (Malawi) and East Africa (Uganda) was conducted based on 82,027 DBLα types found in 4,783 isolates (Table I in Data S3, Figure I in Data S3). Similarly, this spatial study showed that the majority of DBLα types were found at low frequencies with smaller proportions seen at higher frequencies. Comparison of DBLα types and frequencies in the four locations showed conservation of the same upsA DBLα types at moderate-to-high frequencies in all locations (i.e., a highly-frequent DBLα type in Ghana was also found at moderate-to-high frequencies in other analysed locations) (Figure 4). While this was also observed for some highly-frequent DBLα types in the upsB and upsC groups (i.e., non-upsA groups), this study additionally identified some highly-frequent DBLα types in these groups that were present predominantly in a single location, suggesting local selection and preservation of DBLα types in these ups groups (Figure 4, Figure II and III in Data S3). Additionally, as was also observed for isolates in the MRS study, per-isolate frequency profiles in these different locations also consisted of a mix of rare to conserved DBLα types (Figure IV in Data S3). It is worth reminding that these conserved DBLα types make up the minority of all DBLα types in every ups group. An exploration of the relationship between DBLα types and *var* exon 1 sequences revealed that these conserved DBLα types are associated with multiple different *var* exon 1 sequences (Figure V in Data S3), indicating that other parts of the gene were still diversifying even though the DBLα types were maintained in the population. For some of these DBLα types with 1-to-many DBLα-*var* relationships, pairwise nucleotide identity between *var* exon 1 sharing the same DBLα type suggest that some of these *var* sequences could be alleles of a same gene (14). However, the majority of these *var* exon 1 sequences exhibit low shared identity and therefore appear to represent actual different genes (Figure V in Data S3). In a highly-dynamic system where the DBLα domain has been shown *in vitro* to exhibit the highest recombination rate (6), the maintenance of specific DBLα types at high frequencies and through extensive durations could suggest that selection for adaptive advantages.

**Figure 4.**
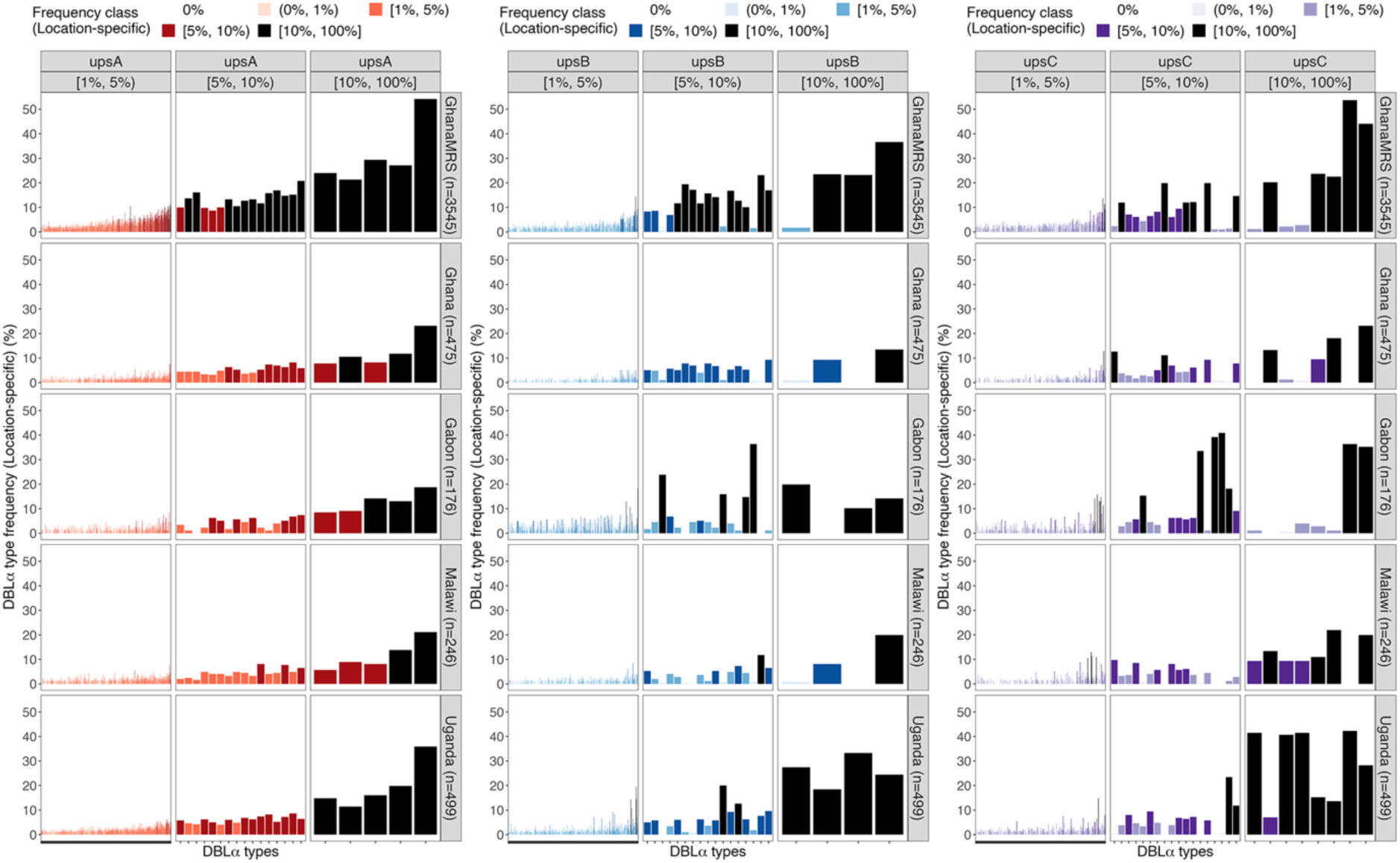
**Conservation of DBLα types and frequencies at local and continent levels [spatial analysis].** Location-specific frequencies of individual DBLα types by ups groups (left to right: upsA, upsB, upsC). Shown here are DBLα types with ≥1% location-averaged frequencies, ordered in increasing frequencies.

### 2.6 Factors driving the conservation of DBLα types remain unknown

The spatial study considered a few possible factors to explain these conserved DBLα types, focusing specifically on 51 and 17 DBLα types in the high and very high location-averaged frequency classes, respectively (i.e., a total of 68 DBLα types with location-averaged frequencies of ≥5%) (Figure 5). Firstly, conservation of *var* genes on specific *P. falciparum* chromosomes 4, 6, 7, and 8 has been previously reported and potentially attributed to selective sweep events associated with antimalarial drug resistance (33). While positional information is unavailable for the DBLα types analysed in this study, sequence alignments show that only six of the 68 highly-frequent DBLα types are homologous to *var* genes on these chromosomes. Furthermore, some of these highly-frequent DBLα types were identified as homologs to the DBLα tags of five *var* genes in *P. praefalciparum*, a Plasmodium species that naturally infects gorillas and is the closest living sister species of *P. falciparum* (41,42). Homologs to the DBLα tag of one *var* gene of another ancestral Laverania species, *P. reichenowi*, was identified but occurring at relatively low-to-moderate frequencies (frequencies range from 0.57% to 2.34%). No homologs to the DBLα tags of *var* genes of *P. gaboni* were identified. Hence, it is clear that while some of these factors can explain the reason a few of these sequences are conserved, the majority of these conserved DBLα types are still unaccounted for.

**Figure 5.**
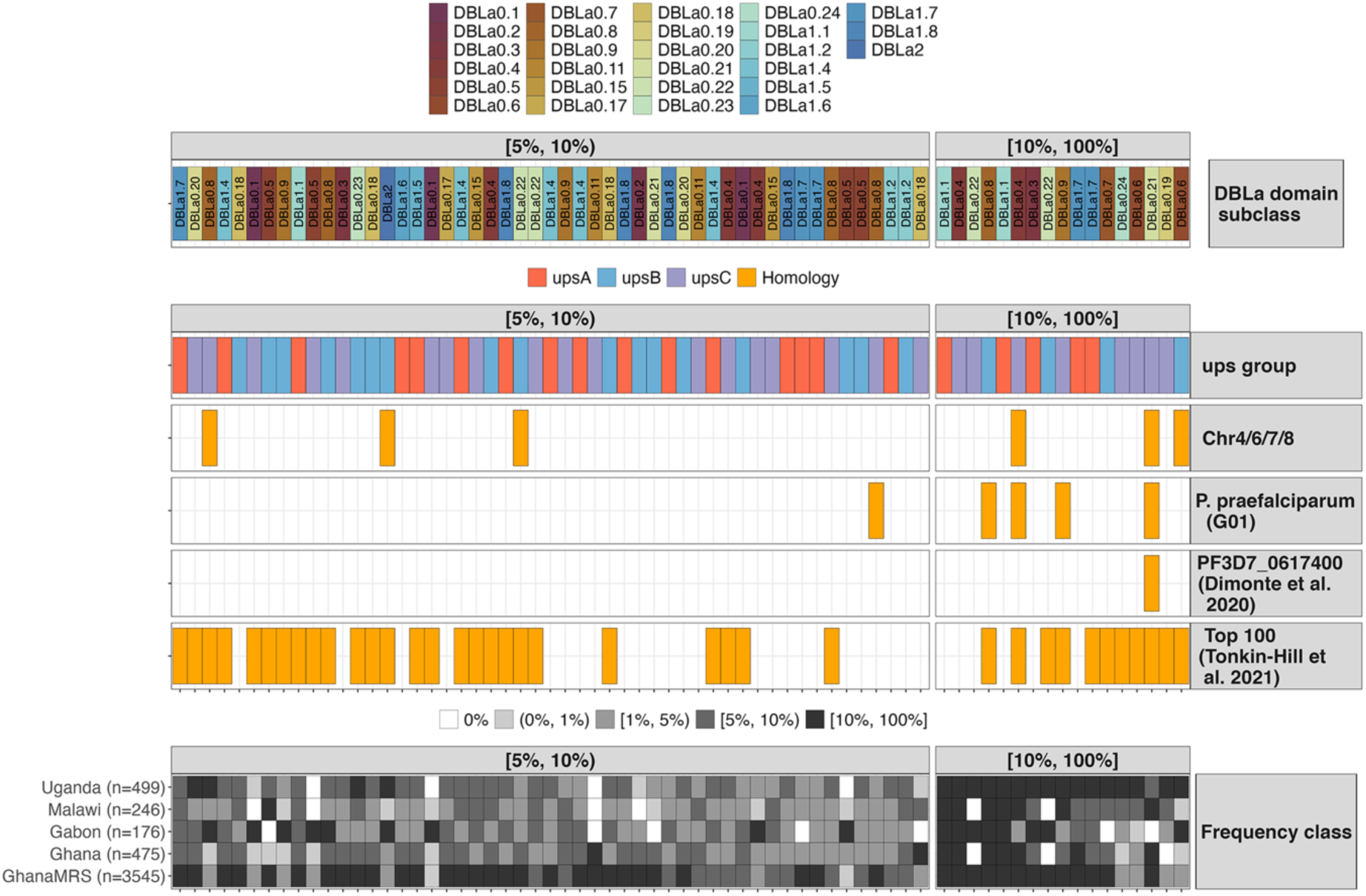
**Annotation of possible factors maintaining conserved DBLα types [spatial analysis].** Vertical panels indicate DBLα types with high or very high frequencies (≥5% location-averaged frequencies) and sequence homology (presence/absence) to published DBLα tag and *var* sequence data.

Homologs to other published globally-conserved DBLα types and *var* gene (PF3D7_0617400) are shown in Figure 5. Tonkin-Hill et al. (35) reported a set of 100 most frequent DBLα types conserved in their analysis of ten countries across diverse global regions in Africa, Asia/Oceania, and South America. Homologs to 84 of these globally-conserved DBLα types were identified in this study, with 26 and 11 of these types found in the two highest location-averaged frequency classes. Furthermore, in the context of general prevalence in the analysed African locations, 30 of these 37 DBLα types were found in all four locations, six in three locations, and only one was found in a single location.

A conserved *P. falciparum var* gene (PF3D7_0617400) was also recently reported in a Gabonese parasite isolate and characterised (34). The homolog to the DBLα tag of this *var* gene was found to occur at high frequencies (ranging from 7.0% to 20.2% in different locations) and present in all locations except, strangely, in Gabon itself. Notably, this conserved PF3D7_0617400 *var* gene is located on chromosome 6, coinciding with the previous reports of haplotypes in linkage disequilibrium on the same chromosome (43,44), though this *var* gene is located outside of this region’s cluster. A possible explanation for the absence of this homolog in the Gabon dataset used in this study may be that the isolates were sampled relatively early in the timeline (year 2000), which precedes the switch to artemisinin (ART)-based combination therapies (ACT) in Africa (45,46), suggesting that the selection for this specific type may have still been in progress and may not yet have risen in frequency to result in observed fixation in the population at the time. Additionally, we also checked for possible conservation of a *P. falciparum var* gene (PF3D7_0809100) that has been reported to be expressed in sporozoites and potentially play a role in hepatocyte infection (47), but homologs to the DBLα tag of this gene were found at only <1% frequency in three locations.

## 3. DISCUSSION

Extensive DBLα type diversity is reported in areas with high malaria transmission, generated by meiotic and mitotic recombination (3–7) with DBLα repertoire diversity driven by frequent outcrossing in the mosquito vector (1,2), such that we would not expect conservation of types. However, a closer inspection of the population structure of DBLα types reveals conservation of DBLα types ***beyond*** sequences found at very high frequencies or within the highest percentiles. Instead, conservation also encompasses types that are seen stable through time and can be found in a population at various frequencies, be it low, moderate, or high. This study observed the conservation of DBLα types in the three major ups groups within a large natural parasite population in a local area in Bongo. Frequencies of these DBLα types at the population level were shown to be temporally stable over at least five years and through wet and dry seasons. In addition to conservation at the continent level, spatial analysis observed local conservation of specific high-frequency upsB and upsC types in individual countries in Africa, despite the high genetic diversity typically reported for these groups. Global analyses such as (35) and (34) would have uncovered the most conserved types and genes globally but could have missed out on much of local signatures of conservation, which this study has shown to exist within different frequency classes.

The key result of this analysis is that the frequency pattern of DBLα types that make up every isolate repertoire not only underlies these local population structures but will maintain them. Looking at individual DBLα types found in every isolate repertoire and the corresponding frequency at which each type occurs in the population, per-isolate frequency profiles revealed that every isolate repertoire consists of a mix of low-, moderate-, and high-frequency types, in proportions consistent across all isolates. This presents a paradox in the population structure of DBLα types, where there is a very high diversity of DBLα types found in a population, but each isolate still maintains a combination of low- to moderate-to high-frequency types in its repertoire. Even more interestingly, this paradoxical structure, both at the level of population and isolate, was observed for types in all three groups of upsA, upsB, and upsC and is maintained despite the expectation of frequent outcrossing in these endemic areas.

In high transmission, despite the high rates of outcrossing and recombination, the consistency in these per-isolate frequency profiles suggest a level of constraint on the modularity of each isolate’s repertoire, i.e., each isolate repertoire must have a combination of common and rare types while maintaining limited overlaps with other isolates in the population overall. This pattern was observed with both DBLα field data and DBLα encoding sequences identified in assembled *var* exon 1 sequences, thereby excluding any biases from genotyping methods. DBLα-*var* relationships revealed that high-frequency DBLα types are likely to be associated with multiple distinct *var* exon 1 sequences (i.e., 1-to-many), though high sequence similarities were estimated for some pairwise *var* sequences, suggesting that a proportion of these are alleles of a same gene. We make clear that this study describes the conservation of DBLα types, not necessarily the conservation of *var* genes. Conservation of *var* genes can be better studied if we can properly define alleles.

Assuming that there are biological advantages conferred by these conserved types, why do we observe stable presence at different frequencies but not fixation of most of these types in the population? Furthermore, why are these per-isolate frequency profiles maintained? One hypothesis relates to balancing selection as a result of co-evolution between the parasite and the human host population it is infecting, which can occur at the local, regional, and continent levels. The role of PfEMP1 in evading recognition by the host innate immune system would select for its variation, and the DBLα domain has been shown to be immunogenic to variant-specific epitopes and serologically recognised in an age-dependent manner (30). On the other hand, its role in virulence, such as the need to bind to specific host endothelial cells for cytoadhesion or blood cell receptors for rosetting, could select for some level of conservation. This tension between the dual different roles, both of which relate to local host genetics, could be creating the observed paradoxical pattern.

Stochastic simulations and network analyses have provided clear evidence for a role of immune selection or negative frequency-dependent selection resulting from specific immune memory, which is a form of balancing selection, in shaping antigenic diversity within natural populations. As antibody-mediated immunity plays a significant role in recognition of PfEMP1 variants, we hypothesise that another possible driver of balancing selection is the arm’s race between the parasite PfEMP1 variants and host HLA class II haplotypes (48–51). Similar to our finding of local signatures of DBLα type conservation against a highly-diverse background, there are also geographic differences in HLA class II alleles across the African continent (52–54). Immune evasion of common local HLA class II alleles could drive DBLα types or *var* genes containing these types to persist through time at stable low-to-moderate frequencies. Alternatively, genetic variation in a parasite’s DBLα or *var* repertoires may have been shaped by underlying differences in host receptors of varying spatial niches and if not, these types could be in linkage disequilibrium with other proximal domains (e.g., CIDR) or genes vital to these roles.

The consistency of these per-isolate frequency profile patterns is striking and suggests that maintaining such frequency profiles within a parasite repertoire is advantageous to the parasite. Having a range of rare to common types may allow malaria parasites to adapt to host factors in order to persist through dynamics and competition within and between hosts. The translational implication of this work suggests that breaking this pattern to what is seen in low transmission i.e., high relatedness of *var* repertoires and clonality could be a target of elimination efforts.

## 4. MATERIALS AND METHODS

### 4.1 Data sources and types

Conservation analyses were performed on a small ∼450bp region of a *var* gene that encodes a portion of the DBLα domain of PfEMP1 (i.e., DBLα tags) (55,56). DBLα tag sequences included in this study were either generated from targeted amplicon sequencing (15–17) or extracted from assembled *var* gene sequences (14,33,40). This made available DBLα tag datasets of varying sizes from Africa and Asia, which were clustered to generate representative DBLα types (see Section 4.2). However, the scope of this study on DBLα conservation was limited to African locations only, with higher transmission, because lower transmission areas may present a different context underlying conservation (e.g., clonality or smaller population sizes). Data in Africa were available from West Africa (Senegal, Gambia, Guinea, Mali, Ghana, Gabon), Central Africa (Congo, Malawi) and East Africa (Uganda, Kenya) (Table I in Data S1, Table I in Data S3). However, most of these African countries were excluded due to limited dataset sizes (number of isolates < 100), resulting in a final analysis from four locations in Africa (i.e., Ghana, Gabon, Malawi, Uganda). Sources and methods that the different studies used to generate these DBLα tag datasets are described in the following subsections.

#### 4.1.1 DBLα tags from targeted amplicon sequencing data

Published DBLα tag datasets from three locations were generated from targeted amplicon sequencing (Table I in Data S1, Table I in Data S3). Amplicon sequencing of DBLα tag sequences involves PCR amplification of a small sequence region encoding the DBLα domain of PfEMP1 with degenerate primers (55,56), followed by high throughput sequencing on either the Illumina MiSeq platform (Ghana, (13,15,39)) or on the 454 sequencing platform (Gabon, (17)), Uganda, (16)). These include sequences from:

I. One area (Bongo) in Ghana: dataset spans seven time points (surveys) from 2012 to 2017 through sampling of asymptomatic individuals through multiple dry and wet seasons.
II. Six areas (Apac, Arua, Jinja, Kanungu, Kyenjojo, Tororo) in Uganda: dataset included sampling of clinical isolates over two years.
III. One area (Bakoumba) in Gabon: dataset included sampling of asymptomatic children in one year.

#### 4.1.2 DBLα tags from assembled *var* gene sequences

Published *var* gene sequences (from isolates in Africa and Asia) were downloaded from the ‘Full Dataset’ published by (33). DBLα tag sequences were identified and extracted from *var* gene sequences (regardless of *var* gene completeness) as described in (14). Briefly, domain annotations provided by (33) were used to extract nucleotide sequences encoding the DBLα domain. These extracted sequences were further translated into the best reading frames and, using *hmmsearch* (57), the resulting amino acid sequences were further searched against positions 189 to 430 of the PFAM profile alignment (PF05424_seed.txt) to identify the ‘tag’ region (domain score cut-off of 60 and ≥100 aligned positions) and to ultimately extract the DBLα tag sequence that would have been amplified with degenerate primers (55,56).

### 4.2 Clustering of DBLα tags into DBLα types

DBLα tags (Africa and Asia) were translated into amino acid sequences and any untranslatable sequences (i.e., stop codons in reading frame) were excluded. The remaining DBLα tags were combined and clustered with *clusterDBLa* (58), using a 96% nucleotide identity threshold (31) to produce representative DBLα types. This also generated a binary matrix detailing the presence/absence matrix of each DBLα type in each isolate.

### 4.3 Classification of DBLα types into domain classes and ups groups

The *classifyDBLα* pipeline (16) was used to classify DBLα types into DBLα domain classes of DBLα0, DBLα1, or DBLα2, in order to confirm that sequences were indeed those encoding the DBLα domain of PfEMP1. In addition, a novel algorithm (***cUps***) described in this study was used to classify DBLα types into the most probable ups group (i.e., upsA, upsB, or upsC), accompanied by assignment probability values. For each DBLα type, ups groups were assigned according to the prediction with the highest assignment probability. We describe this novel classification algorithm below as well as in Data S2). An implementation of the algorithm is available at https://github.com/qianfeng2/cUps.

Through the alignment and clustering of 2kb sequences upstream of *var* genes, followed by the classification *var* genes into ups groups by Neighbour-joining (NJ) and Markov clustering (MCL) methods (trees available in Data S2), a reference dataset of DBLα tag sequences was generated from 846 *var* genes from 16 P. falciparum genomes ((59) and NCBI). We begin with this reference database of DBLα tag sequences with ups groups and DBLα subclasses known. For each category (ups group/DBLα subclass combination), we align the reference sequences in the category using Clustal Omega v1.2.4 (60), then fit a profile hidden Markov model (61) using HMMER v3.2.1 (57) with default settings.

For a given query sequence (representing a DBLα type), we calculate the likelihood of the query sequence being drawn from the profile HMM of each category, using the forward algorithm. The posterior probability for each category is then calculated using Bayes’ Theorem, with the prior probabilities of each category calculated from the reference database. Summing over DBLα subclasses gives the posterior probability for each ups group (i.e., assignment probability). The query sequence can be classified to the ups group with the highest assignment probability. Although we do not do so in this paper, a threshold may optionally be applied, so that sequences with highest assignment probability below the threshold are categorised as ‘unclassified’. Alternatively, a summary statistic may weight each ups group by the assignment probability. This method is described in much more detail, with verification (Feng, submitted).

### 4.4 Exclusion of DBLα types, isolates, and populations from the final DBLα type dataset

Only the DBLα types that were successfully classified into a DBLα domain class (i.e., DBLα0, DBLα1, or DBLα2) were retained in the final dataset. Subsequently, isolates with < 20 DBLα types were also removed from dataset to ensure robust analyses downstream (Table I in Data S1, Table I in Data S3). Specifically for the time-series dataset from the Malaria Reservoir Study (MRS) in Bongo, Ghana (13,15,39), submicroscopic or symptomatic isolates were additionally excluded from the dataset. Further, using *blastn* (≥96% nucleotide identity, ≥95% query coverage) (62), DBLα types with homology to isolate-transcendent *var1*, *var2csa*, and *var3* sequences (sequences from (22,33)) were excluded to remove putative DBLα types previously reported as isolate-transcendent (20,21). Finally, given that frequency classes and profiles were calculated based on proportional frequencies, only locations with datasets of ≥100 isolates were retained. This resulted in the exclusion of six African countries from this study (“*” in Table I in Data S3).

### 4.5 Genetic similarity between pairwise isolate repertoires

The pairwise type sharing metric (PTS) (31) is used to estimate the overlap between pairwise isolate repertoires (e.g., isolates *i* & *j*). Specifically:

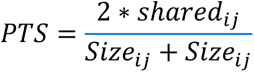

where *shared_ij_* is the number of shared DBLα types between repertoires of isolates *i* and *j*, and *Size_i_* and *Size_j_* are the total number of DBLα types (i.e., repertoire sizes) of isolates *i* and *j*, respectively. A value of 0 indicates the absence of sharing between two isolates whereas a value of 1 indicates completely identical isolate repertoires.

### 4.6 Calculation of DBLα type frequencies and assignments into frequency classes

Depending on the analysis, a population can be the collection of isolates sampled at a specific survey or time point in the time-series analyses (i.e., by year or survey in the MRS data) or the collection of isolates sampled from a specific region or location/country in the spatial analyses. Raw frequencies of DBLα types were defined at the survey or location level in counts (i.e., number of isolates with a particular DBLα type in each survey or location). Raw (count) frequencies were converted into proportional frequencies through division of count frequencies by the total number of isolates at a corresponding time point or location, leading to “survey-specific frequencies” or “location-specific frequencies”. Subsequently, these frequencies were further categorised into frequency classes of 0%, ***low*** (0%, 1%), ***moderate*** [1%, 5%), ***high*** [5%, 10%), and ***very high*** [10%, 100%].

Given the substantial differences in dataset sizes across surveys or locations (e.g., 499 isolates for Uganda *versus* 176 isolates for Gabon), simply summing isolates across datasets of multiple surveys or locations would bias total frequencies to reflect those of larger datasets. Hence, averaged frequencies were used instead as a means to normalise total frequencies by isolate counts in each survey or location. For example, a DBLα type found in 10 out of 100 isolates for location A and 10 out of 500 isolates for location B would be reported to have 10% and 2% frequencies for locations A and B, respectively. This would yield a crude total frequency of 3.33% (20 of 600 isolates), which is more reflective of the frequency observed in location B even though the DBLα type was found at relatively high frequency at location A. In this instance, with normalisation, an averaged frequency of 6% would be estimated (12 of 200 isolates), reducing the bias towards larger dataset sizes. Given the focus of this study on conserved DBLα types, this normalisation method provides a less biased approach in identifying DBLα types that are found at high frequencies but not necessarily uniformly across all datasets.

### 4.7 Determination of DBLα-*var* relationships

For two locations (Ghana and Malawi), *var* gene sequences were available from assemblies performed by (33). DBLα-*var* relationships were determined using complete *var* exon 1 sequences that are bounded by an N-terminal segment (NTS) and a transmembrane region (TM) on the 5’ and 3’ ends of exon 1, respectively (14). Briefly, using *vsearch* (63), DBLα types were globally aligned to *var* exon 1 sequences from the same location (e.g., Malawi DBLα types to Malawi *var* exon 1). Given that DBLα types were generated from clustering at a 96% nucleotide identity threshold, these types were aligned to *var* exon 1 sequences at the same threshold of 96% identity, calculated over the alignment length and excluding terminal gaps *(--iddef 2*). The relationship between a DBLα type and distinct *var* exon 1 was determined based on the number of unique *var* exon 1 sequences sharing a same DBLα type (e.g., a 1-to-*n* DBLα-*var* relationship is defined as a DBLα type found in *n* unique *var* exon 1).

For each group of *var* exon 1 that share a same DBLα type, an all *vs* all sequence alignment of *var* exon 1 sequences in the group was performed using the *allpairs_global* option within *vsearch* (63) and set to include all pairwise alignments *(--acceptall*). Pairwise nucleotide identities were estimated based on calculations over whole alignment lengths, including terminal gaps (*--iddef 1*), to account for differences in pairs of *var* exon 1 of variable lengths.

### 4.8 Search for homology to other DBLα types or *var* genes

#### 4.8.1 *Var* genes in association with selective sweeps on select chromosomes

Published work reported conserved *var* genes on chromosomes 4, 6, 7 and 8 associated with selective sweep events, potentially due to drug resistance or other factors. Accession numbers of these conserved genes were obtained from the author (33,59) and used as reference. Using *blastn* (62), DBLα types were searched against these reference sequences and hits from alignments were reported (≥96% nucleotide identity, ≥95% query coverage).

#### 4.8.2 *Var* genes in primate *Plasmodium* species

*Var* genes from three *Plasmodium* species, *P. praefalciparum*, *P. reichenowi* and *P. gaboni*, were downloaded from PlasmoDB (41) and used as reference. Using *blastn* (62), DBLα types were searched against these reference sequences and hits from alignments were reported (≥96% nucleotide identity, ≥95% query coverage).

#### 4.8.3 Globally-conserved DBLα types or *var* genes

The 100 most frequent DBLα sequences reported in the global analysis by (35) was used as reference. Using *blastn* (62), DBLα types were searched against these reference sequences and hits from alignments were reported (≥96% nucleotide identity, ≥95% query coverage). The same search parameters and thresholds were applied in searching for homologs to *var* gene sequence PF3D7_0617400, a conserved 3D7 *var* gene reported by (34). Homologs to *var* gene sequence PF3D7_0809100, shown by (47) to be expressed at the sporozoite stage, were also searched for.

## ACKNOWLEDGMENTS

This publication uses *var* gene data assembled from data generated from the MalariaGEN *Plasmodium falciparum* Community Project. This research was supported by The University of Melbourne’s Research Computing Services and the Petascale Campus Initiative.

