## Supplementary material for "A paradoxical population structure of *var* DBLα types in Africa": Data S1

### Data S1. Study of DBLα conservation in Bongo, Ghana

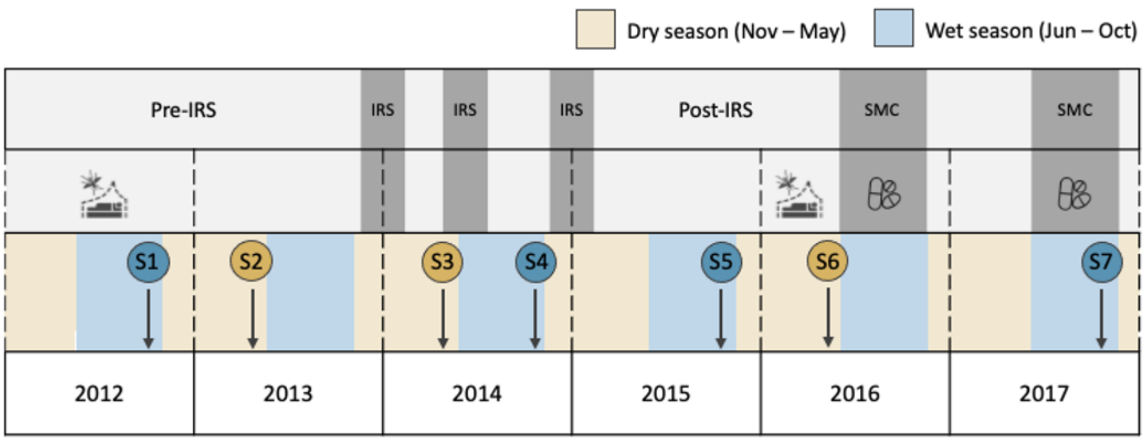


**Figure I. Schedule of seven age-stratified cross-sectional surveys in the Malaria Reservoir Study (MRS) conducted in Bongo, Ghana.** The study area is typically characterised by high seasonal malaria transmission (prevalence ≥ 35% at baseline (i.e., S1)). Wet- and dry-season surveys (S1 to S7) are represented by the blue and yellow coloured circles, respectively. Interventions (i.e., bed nets, indoor residual spraying (IRS), and seasonal malaria chemoprevention (SMC)) are also shown.

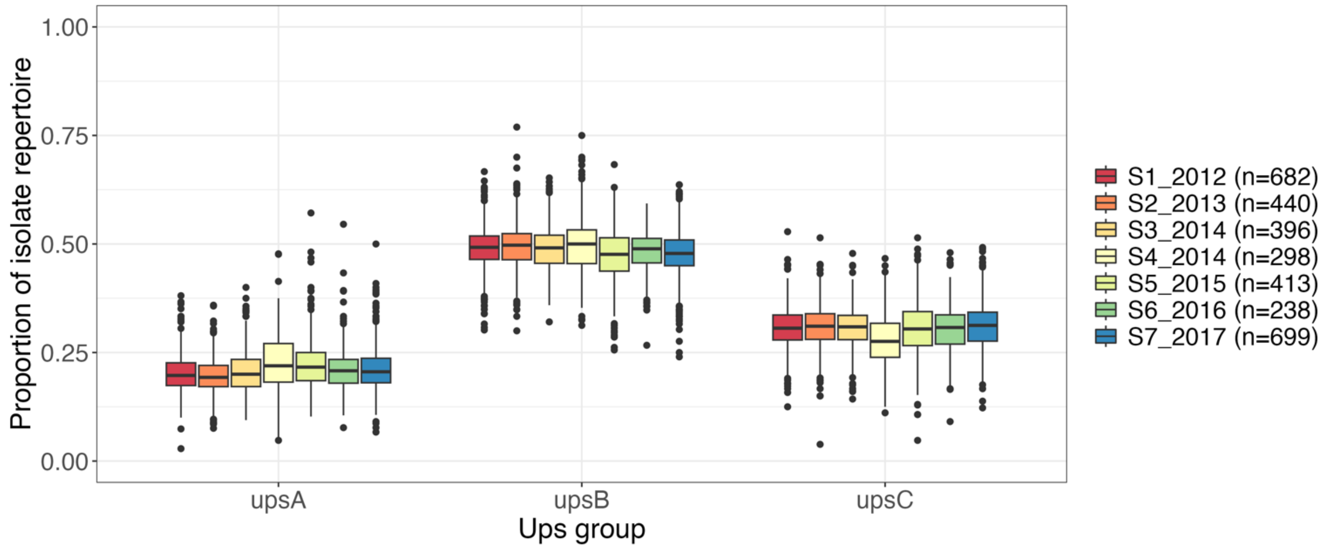


**Figure II. Proportions of upsA, upsB, and upsC DBLα types per isolate repertoire [Malaria Reservoir Study (MRS)].**

**Table I. Details on participants, microscopic characteristics, and DBLα types in the Malaria Reservoir Study (MRS) surveys**

| Parasitological parameters | Survey 1 (S1) | Survey 2 (S2) | Survey 3 (S3) | Survey 4 (S4) | Survey 5 (S5) | Survey 6 (S6) | Survey 7 (S7) |
|---|---|---|---|---|---|---|---|
| | Oct (2012) | Jun (2013) | Jun (2014) | Oct (2014) | Oct (2015) | Jun (2016) | Oct (2017) |
| **Number of participants**[a] | 1,923 | 1,902 | 1822 | 1,866 | 2,022 | 2,091 | 1,915 |
| **Isolates with microscopically positive (prevalence**[b]**)** | 808 (42.0) | 513 (27.5) | 535 (29.4) | 430 (23.0) | 545 (27.0) | 272 (13.0) | 789 (41.2) |
| **Isolates with > 0 DBLα data**[c] | 742 (91.8) | 468 (91.2) | 488 (91.2) | 386 (89.8) | 508 (93.2) | 261 (96.0) | 759 (96.2) |
| **Isolates with ≥ 20 DBLα data**[c] | 682 (84.4) | 440 (85.8) | 396 (74.0) | 298 (69.3) | 413 (75.8) | 238 (87.5) | 699 (88.6) |
| **Total DBLα types** | | | | 62,168 | | | |
| **upsA** | | | | 3,370 (5.4) | | | |
| **upsB** | | | | 35,215 (56.6) | | | |
| **upsC** | | | | 23,583 (37.9) | | | |
| **Number of DBLα types per survey**[d] | 33,746 | 26,627 | 24,059 | 15,749 | 18,686 | 17,255 | 28,167 |
| **upsA** | 2,398 (7.1) | 2,012 (7.6) | 1,958 (8.1) | 1,602 (10.2) | 1,795 (9.6) | 1,627 (9.4) | 2,252 (8.0) |
| **upsB** | 19,353 (57.3) | 15,271 (57.4) | 13,733 (57.1) | 9,137 (58.0) | 10,415 (55.7) | 9,633 (55.8) | 15,922 (56.5) |
| **upsC** | 11,995 (35.5) | 9,344 (35.1) | 8,368 (34.8) | 5,010 (31.8) | 6,476 (34.7) | 5,995 (34.7) | 9,993 (35.5) |
| **Number of DBLα types per isolate** | | | | | | | |
| **Mean** | 167.3 | 163.4 | 156.7 | 108.7 | 99.5 | 148 | 127.6 |
| **Median** | 141.5 | 140 | 116.5 | 78 | 76 | 113.5 | 105 |
| **Min** | 20 | 20 | 21 | 20 | 20 | 21 | 20 |
| **Max** | 904 | 594 | 719 | 607 | 619 | 658 | 549 |

[a] Number of participants surveyed that were analysed by microscopy.

[b] Data reflect the number (% (n/N)) of participants sampled that were microscopically positive for an asymptomatic *P. falciparum* infection (including mixed *P. falciparum* infections) relative to the number of participants surveyed in the total population or by the age groups presented.

[c] Data reflect the number (% (n/N)) of microscopic *P. falciparum* isolates that had DBLα sequence data relative the number of participants surveyed that were microscopically positive for *P. falciparum* (including mixed *P. falciparum* infections) in the total population. Isolates that had low DNA and/or sequencing quality (i.e., < 20 DBLα types) were removed from downstream analyses (see Methods for additional details).

[d] Data reflect DBLα types that have been confirmed through successful categorization into DBLα domain classes.

#### Table II. Classification of DBLα types in the Malaria Reservoir Study (MRS) surveys into ups groups and frequency classes

| Malaria Reservoir Study (MRS) surveys | S1 | S2 | S3 | S4 | S5 | S6 | S7 |
|---|---|---|---|---|---|---|---|
| | Oct (2012) | Jun (2013) | Jun (2014) | Oct (2014) | Oct (2015) | Jun (2016) | Oct (2017) |
| **Number of DBLα types per survey** [a] | 33,746 | 26,627 | 24,059 | 15,749 | 18,686 | 17,255 | 28,167 |
| upsA [b] | 2,398 (7.1) | 2,012 (7.6) | 1,958 (8.1) | 1,602 (10.2) | 1,795 (9.6) | 1,627 (9.4) | 2,252 (8.0) |
| (0%, 1%) [c] | 1599 (66.7) | 1282 (63.7) | 1168 (59.7) | 933 (58.2) | 1270 (70.8) | 913 (56.1) | 1542 (68.5) |
| [1%, 5%) [c] | 662 (27.6) | 575 (29.1) | 650 (33.2) | 583 (36.4) | 462 (25.7) | 574 (35.3) | 619 (27.5) |
| [5%, 10%) [c] | 100 (4.2) | 106 (5.3) | 105 (5.4) | 72 (4.5) | 51 (2.8) | 105 (6.5) | 73 (3.2) |
| [10%, 100%) [c] | 37 (1.5) | 39 (1.9) | 35 (1.8) | 14 (0.9) | 12 (0.7) | 35 (2.2) | 18 (0.8) |
| upsB [b] | 19,353 (57.3) | 15,271 (57.4) | 13,733 (57.1) | 9,137 (58.0) | 10,415 (55.7) | 9,633 (55.8) | 15,922 (56.5) |
| (0%, 1%) [c] | 17674 (91.3) | 13753 (90.1) | 11816 (86.0) | 7801 (85.4) | 9876 (94.8) | 8140 (84.5) | 14799 (92.9) |
| [1%, 5%) [c] | 1625 (8.4) | 1458 (9.5) | 1864 (13.6) | 1306 (14.3) | 513 (4.9) | 1441 (15.0) | 1087 (6.8) |
| [5%, 10%) [c] | 35 (0.2) | 42 (0.3) | 35 (0.3) | 18 (0.2) | 17 (0.2) | 38 (0.4) | 21 (0.1) |
| [10%, 100%) [c] | 19 (0.1) | 18 (0.1) | 18 (0.1) | 12 (0.1) | 9 (0.1) | 14 (0.1) | 15 (0.1) |
| upsC [b] | 11,995 (35.5) | 9,344 (35.1) | 8,368 (34.8) | 5,010 (31.8) | 6,476 (34.7) | 5,995 (34.7) | 9,993 (35.5) |
| (0%, 1%) [c] | 11033 (92.0) | 8463 (90.6) | 7247 (86.6) | 4321 (86.2) | 6111 (94.4) | 5140 (85.7) | 9269 (92.8) |
| [1%, 5%) [c] | 917 (7.6) | 832 (8.9) | 1075 (12.8) | 665 (13.3) | 341 (5.3) | 815 (13.6) | 690 (6.9) |
| [5%, 10%) [c] | 28 (0.2) | 30 (0.3) | 30 (0.4) | 14 (0.3) | 12 (0.2) | 25 (0.4) | 21 (0.2) |
| [10%, 100%) [c] | 17 (0.1) | 19 (0.2) | 16 (0.2) | 10 (0.2) | 12 (0.2) | 15 (0.3) | 13 (0.1) |

[a] Data reflect DBLα types that have been confirmed through successful categorisation into DBLα domain classes and ups groups.

[b] Data reflects the number (% (n/N)) of DBLα types relative to the number in a survey (i.e., "Number of DBLα types per survey").

[c] Data reflects the number (% (n/N)) of DBLα types relative to the number in a survey and ups group (i.e., "upsA", "upsB", or "upsC").

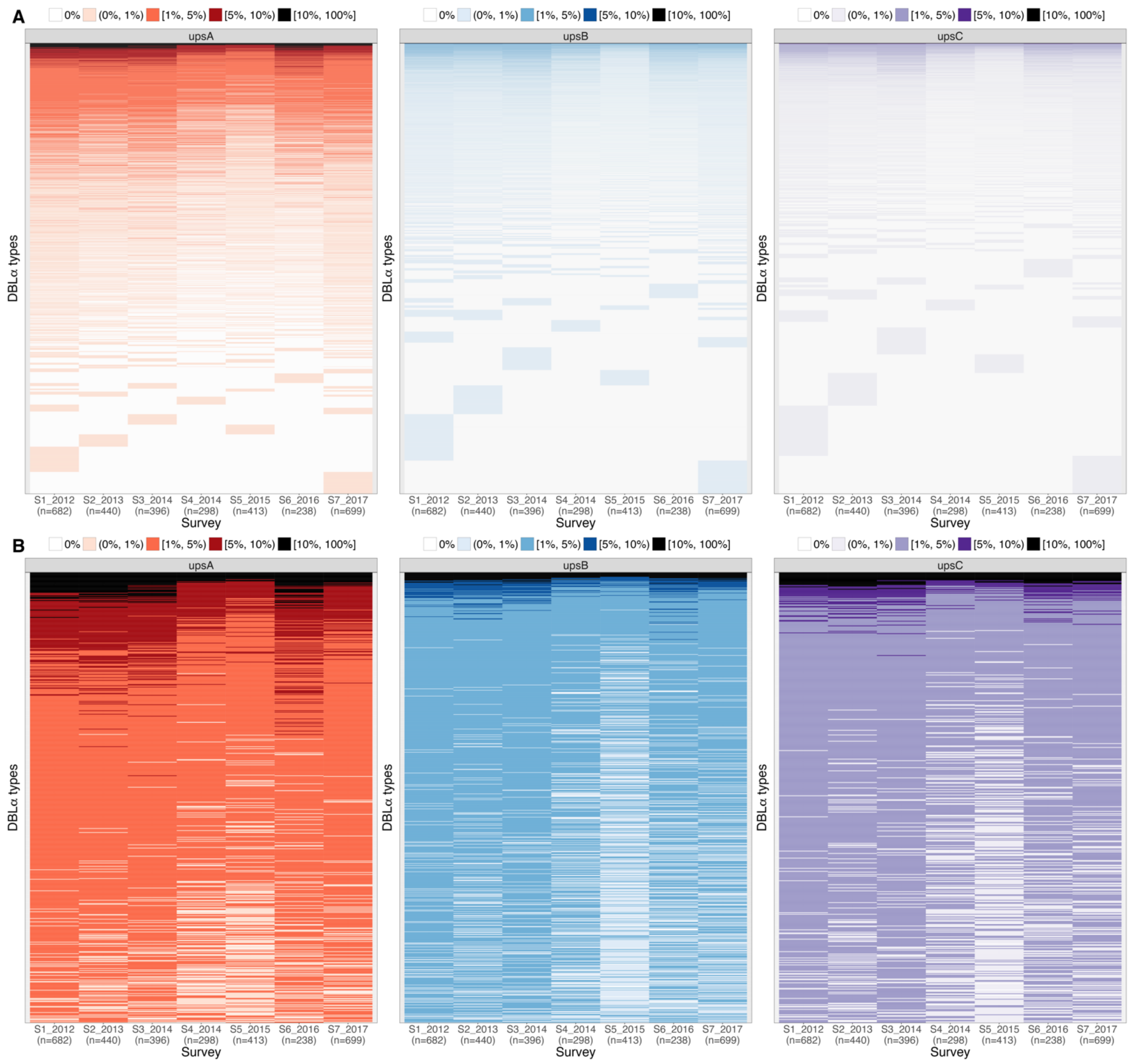


**Figure III. DBLα types and survey-specific frequency classes [Malaria Reservoir Study (MRS)].** Plots show (A) all DBLα types and (B) DBLα types at ≥1% frequencies (survey-averaged frequencies). Rows represent individual DBLα type in each ups group, ordered in decreasing survey-averaged frequencies. Colours reflect the ups groups and survey-specific frequency classes of DBLα types.

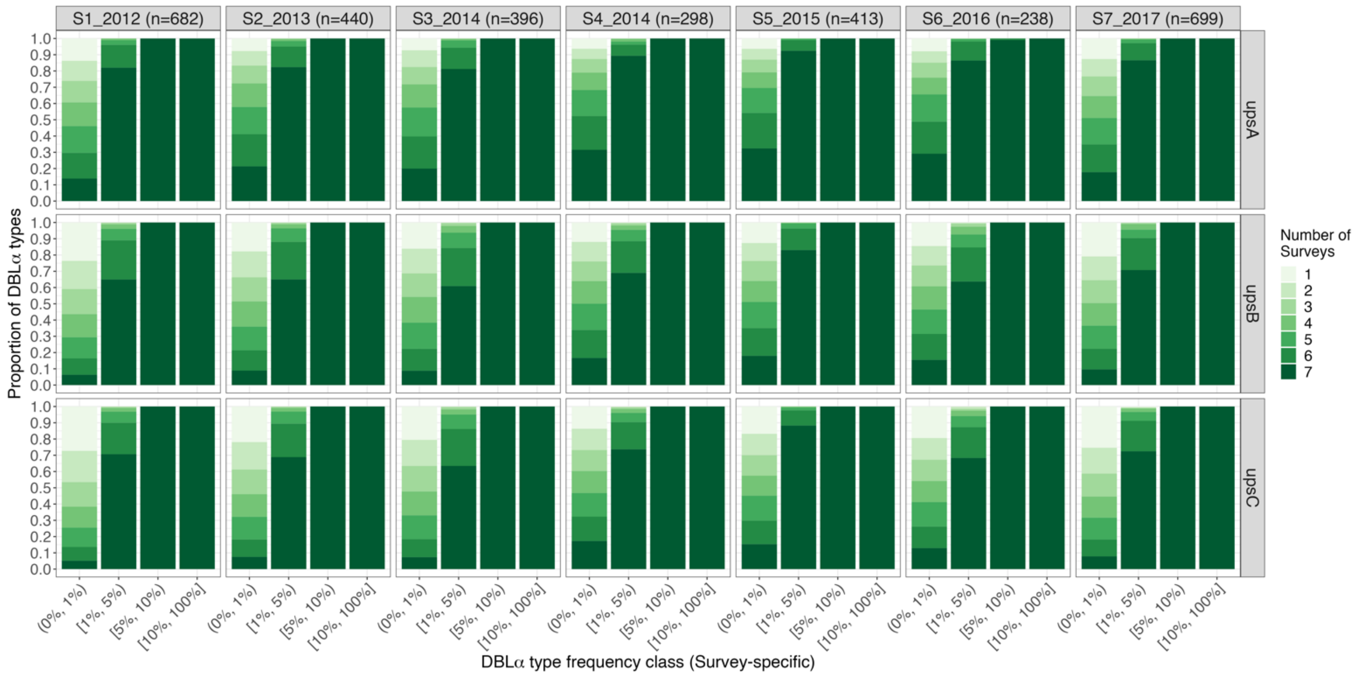


**Figure IV. Number of surveys DBLα types were observed in, showing that DBLα types found at ≥1% survey-specific frequencies were seen to also persist through most surveys [Malaria Reservoir Study (MRS)].**

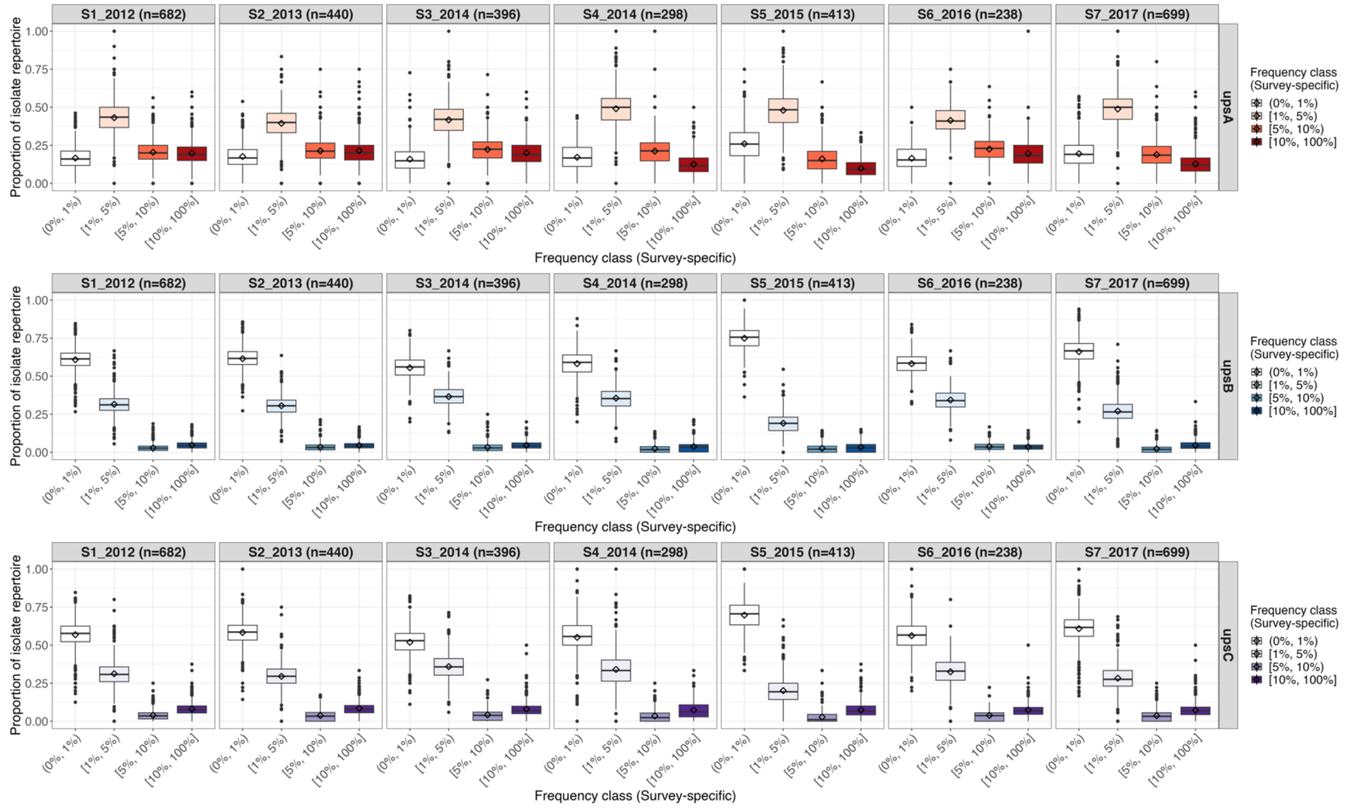


**Figure V. Proportions of survey-specific frequency classes within per-isolate frequency profiles [Malaria Reservoir Study (MRS)].** Box plots show distributions of per-isolate proportions for the different survey-specific frequency classes, by ups groups and surveys. Boxes represent the interquartile range (IQR, i.e. 25th to 75th percentiles). Horizontal lines and diamonds in each box plot represent median and mean values, respectively. Whiskers span 1.5 times the IQR above the upper quartile and below the lower quartile.

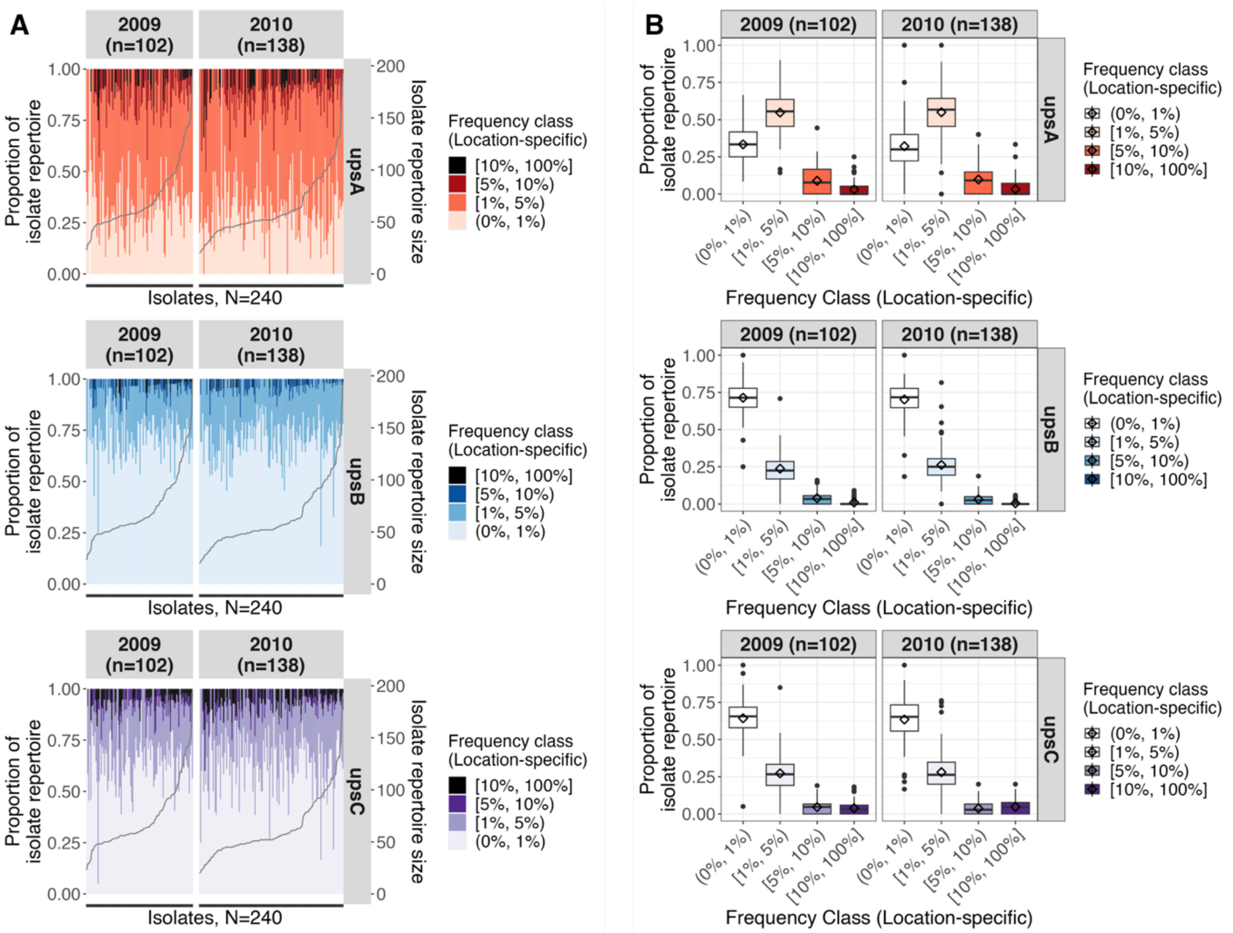


**Figure VI. Per-isolate frequency profiles show the composition of time-specific frequency classes in every isolate repertoire, using DBLα tag sequences extracted from assembled *var* gene data found in Navrongo, Ghana**. (A) Vertical bars represent individual isolates, showing the composition of frequency classes in each isolate (left y-axis) by ups group (horizontal panels). Isolates are sorted by isolate repertoire size in increasing order, with isolate repertoire sizes indicated by the grey line (right y-axis). (B) Box plots show distributions of per-isolate proportions for the different survey-specific frequency classes, by ups groups and surveys. Boxes represent the interquartile range (IQR). Horizontal lines and diamonds in each box plot represent median and mean values, respectively. Whiskers span 1.5 times the IQR above the upper quartile and below the lower quartile.

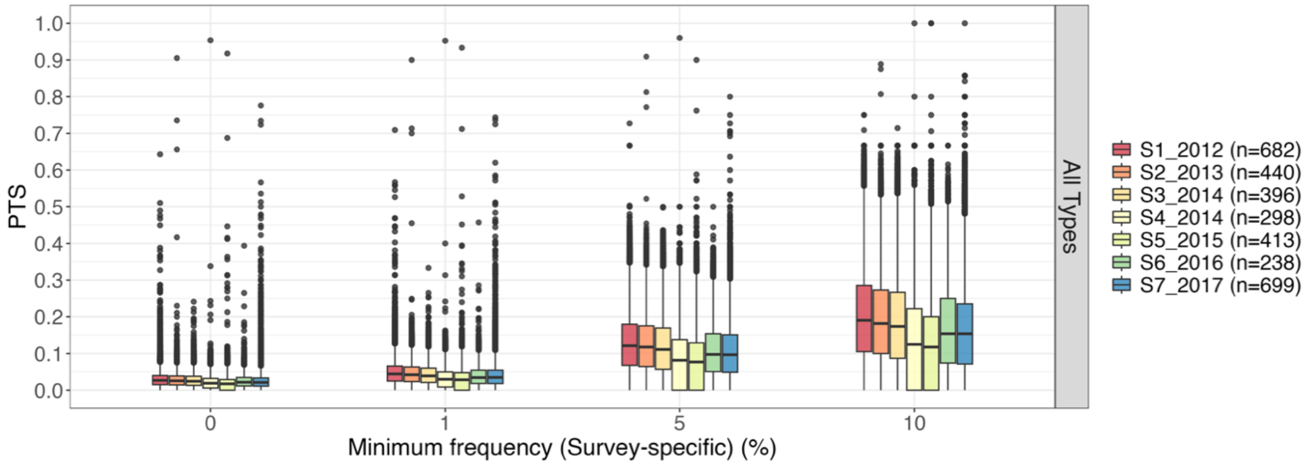


**Figure VII. Similarity of isolate repertoires by pairwise type sharing (PTS), based on DBLα types occurring at minimum survey-specific frequencies of 0%, 1%, 5%, and 10% [Malaria Reservoir Study (MRS)].**
