## Supplementary material for "A paradoxical population structure of *var* DBLα types in Africa": Data S2

### Data S2. A probabilistic method for classifying the malaria *var* genes into ups groups (*cUps*)

**The *cUps* algorithm can be found at https://github.com/qianfeng2/cUps**

##### Data S2.1 Creating the reference dataset

Based on methods described by (1), *var* gene sequences were extracted from publicly-available *P. falciparum* genomes and, using their 2kb upstream sequences, assigned to ups groups by Neighbor joining (NJ) and Markov clustering (MCL). Assigned to ups groups, the corresponding DBLα tag sequences were used as reference dataset for the ***cUps*** algorithm mentioned in in Materials and Methods (main paper, section 4.3).

To elaborate, genome assembly and annotation files of a total of 18 *Plasmodium falciparum* strains were downloaded from PlasmoDB release 51 (16 strains: 3D7, 7G8, CD01, Dd2, GA01, GB4, GN01, HB3, IT, KE01, KH01, KH02, ML01, SD01, SN01, and TG01) and NCBI (2 strains: IGH-CR14, RAJ116), described in (1,2) to obtain *var* gene sequences and their corresponding 2kb upstream sequences. These strains were selected because they were either long-read assemblies with characterised subtelomeric regions (2) and/or because their *var* genes have been previously described (1). For each genome, GFF-format annotation files were searched for genes assigned with as "PfEMP1", excluding sequences of pseudogenes, exon 2, or acidic terminal segment (ATS) regions (i.e., those containing keywords such as "pseudogene", "exon 2", or "acidic" in their descriptions). Nucleotide and amino acid sequences for the remaining PfEMP1 genes were extracted from coding sequence and protein files that were downloaded from the respective sources. The VarDom 1.0 server (1) as used to predict PfEMP1 domains. To properly identify 2kb upstream sequences, only genes with a NTS domain predicted in the VarDom annotation were included as this domain typically marks the N-terminus of PfEMP1 proteins (1). Bedtools (3) was used to extract genomic sequences 2000 bases upstream from the start of each *var* gene. Using the *hmmscan* in HMMER v3.3.1 (4), DBLα tag sequences were identified based on homology to positions 189 to 430 of the Pfam domain (PF05424). Using *classifyDBLa* v1.0 (5), DBLα tag sequences were classified into different DBLα/pam domain classes (i.e. DBLα0, DBLα1, DBLα2, or DBLpam). Detailed information on the method of extracting DBLα tags and classification into upsA/non-upsA groups is described in (6) and (5), respectively.

Similar to described by (1), multiple sequence alignment of the 2kb upstream sequences was performed with MAFFT v7.471 (7) using the L-INS-i algorithm. ClustalW v2.1 (8) was used to generate a bootstrapped (n=1000) neighbor-joining tree. Upstream sequences were also clustered using the Markov clustering algorithm, following the parameters implemented in (1). This included firstly an all-vs-all pairwise alignment of upstream sequences was performed with *blastn* v2.10.1 (9), followed by clustering with the MCL v14-137 algorithm (10). The same inflation parameters (varied in steps of 0.2 from 1.2 to 5.0) and resource scheme 7 (most accurate) was used, as reported in (1). A consensus of all clustering trees was generated by Majority Rule (with extensions) using Phylip v3.697 (11). Both the NJ and MCL trees were visualised with iTOL v6 (12), alongside assignment of ups groups by Rask et al 2017 where available, DBLα/pam domain classes, and PfEMP1 domain structure (Figure I and Figure II in this document). As ML01 and TG01 were identified as mixed infections in (2), these isolates were excluded from the final reference dataset. Based on results from the NJ and MCL methods, *var* genes with discordant assignments were excluded from the reference dataset. This resulted in a final consensus ups classification of 846 *var* genes from 16 *P. falciparum* genomes, from which the DBLα tag was included in the reference dataset.

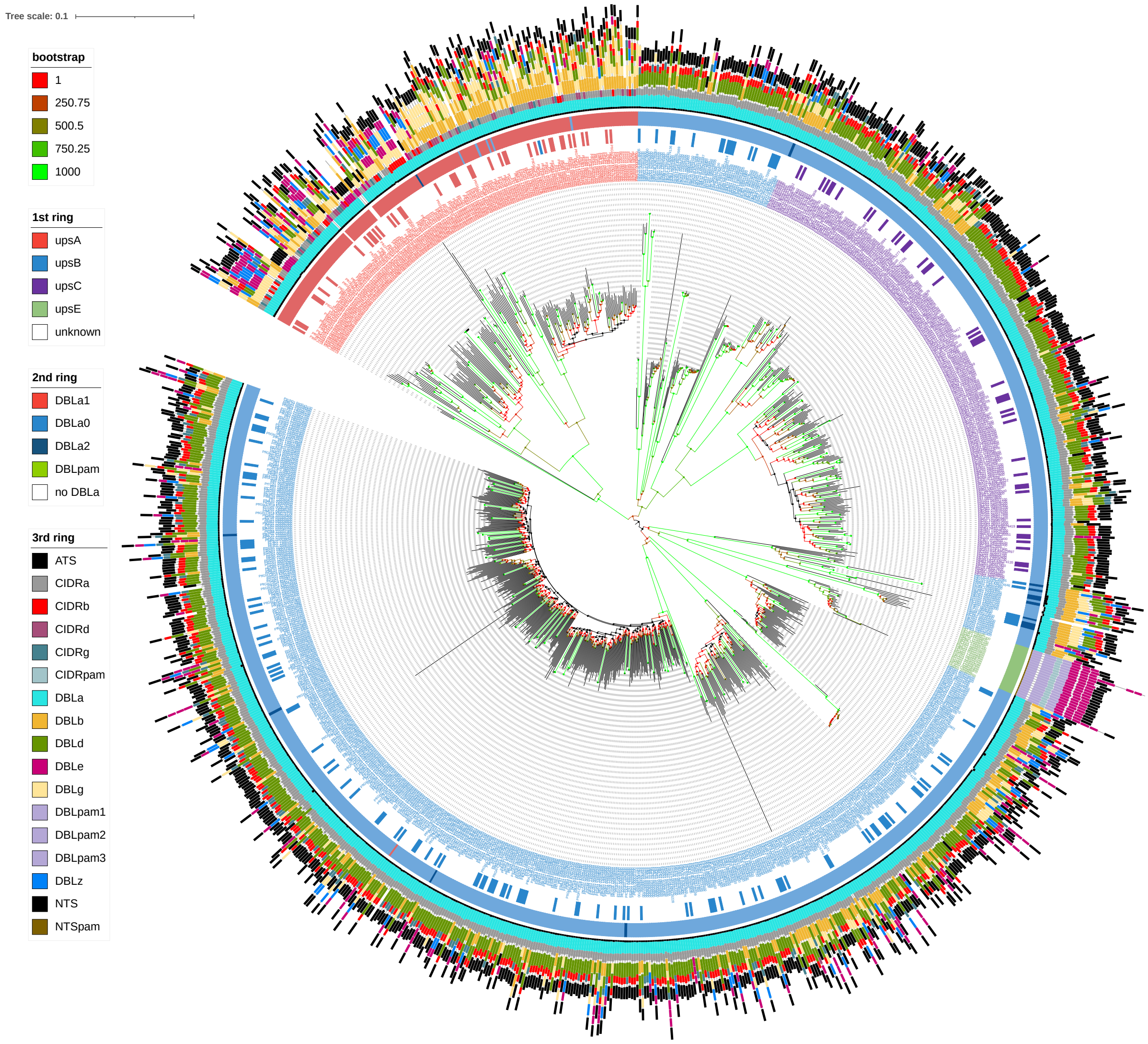


**Figure I. Classification of *var* genes into ups groups by Neighbour-joining, using 2kb upstream sequences.** Tips (leaves) of the tree represent each *var* gene ID, coloured by the ups classification based on clustering shown in this tree. The inner-most (first) ring shows classification of *var* genes into upsA/B/C/E groups by (1) where available. The second ring indicates DBLα/pam domain classes, classified using *classifyDBLa* (5). The outer-most ring shows the PfEMP1 domain composition and structure.

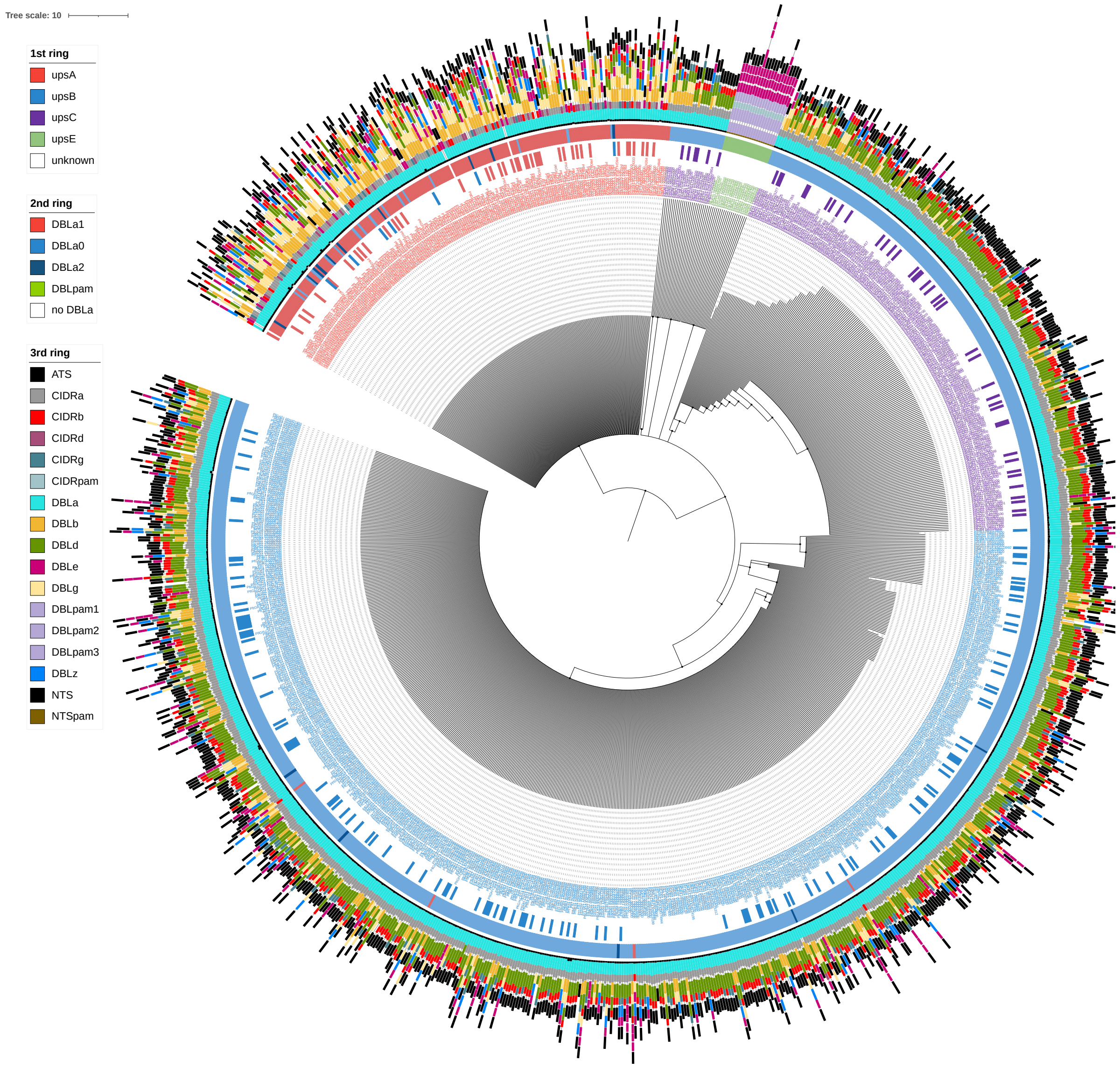


**Figure II. Classification of *var* genes into ups groups by Markov clustering algorithm, using 2kb upstream sequences.** Tips (leaves) of the tree represent each *var* gene ID, coloured by the ups classification based on clustering shown in this tree. The inner-most (first) ring shows classification of *var* genes into upsA/B/C/E groups by (1) where available. The second ring indicates DBLα/pam domain classes, classified using *classifyDBLa* (5). The outer-most ring shows the PfEMP1 domain composition and structure.

**Data S2.2 Accuracy of ups classification algorithm**
**Data S2.2.1 Performance on reference database**

We evaluated the accuracy of our novel ups classification algorithm via leave-one-out cross-validation on the reference database, consisting of 846 sequences (148 upsA, 502 upsB, 196 upsC; see Section Data S2.1 above for further details).

In Table I below, we show the confusion matrix for our new method, and in Table II below we assess the accuracy of our method using two standard measures: sensitivity (the proportion of each group that is correctly predicted) and precision (the proportion of predictions for each group that is correct). It is clear that upsA can be classified very accurately; in contrast, although the method retains good predictive power for upsB and upsC, the accuracy is considerably lower. This is consistent with existing methods, which are unable to distinguish B from C at all.

**Table I. Confusion matrix for ups classification algorithm using leave-one-out cross-validation on the reference database.**

| | | **Real** | | | |
|---|---|---|---|---|---|
| | | upsA | upsB | upsC | **Total** |
| **Predicted** | upsA | **145** | 1 | 1 | 147 |
| | upsB | 3 | **350** | 54 | 407 |
| | upsC | 0 | 151 | **141** | 292 |
| | Total | 148 | 502 | 196 | 846 |

**Table II. Accuracy measures for ups classification algorithm using leave-one-out cross-validation on the reference database.**

| | upsA | upsB | upsC |
|---|---|---|---|
| Sensitivity | 98.0% | 69.7% | 71.9% |
| Precision | 98.6% | 86.0% | 48.3% |

**Data S2.2.2 Comparison to existing pipeline**

We compared our method to the existing pipeline (1,5). This pipeline first determines the DBLα subclass of the query tag sequence, and classifies DBLα1 sequences to the upsA group and other sequences to the upsB/C group. This method does not distinguish between the upsB and C groups (we know of no method in the literature that does).

Similarly to our method, the existing pipeline can very accurately classify a sequence to upsA or non-upsA when using leave-one-out cross-validation on the reference database (4 upsA and 1 ups B/C misclassified).

The 3 upsA sequences that are misclassified with our method (identifiers "PfGB4_000018400", "PfGN01_100005800", "PfHB3_120045300") are distinct from the 4 upsA sequences that are misclassified with the existing pipeline ("PfIGH-CR14_KNG75243", "PfKH02_030030250", "PfKH01_130076800", "PfGA01_010005600"). In contrast, both methods misclassify the same upsB sequence ("PfGN01_020028200") to upsA, suggesting that the labelling of this sequence in the reference database may be unreliable. Our method additionally misclassifies an upsC sequence ("PfIGH-CR14_KNG77255") to upsA.

In addition to the ability to distinguish the upsB and upsC groups, our method also provides meaningful classification (posterior) probabilities that can be exploited for further information. This enables a more detailed comparison of the accuracy of the two methods by considering the output as the classification probabilities (taken as a vector of length 3). For the existing pipeline, we consider a non-upsA classification to have a probability of 0.5 for both upsB and upsC, while an upsA classification has a probability of 1 for upsA as expected.

We compared the methods using three measures: overall accuracy (OA) and kappa coefficient (for definitions see (13)), and root mean square error (RMSE). We show the results in Table III below. Here, we see that our method has significantly better OA and kappa, and similar RMSE to the existing pipeline.

**Table III. Accuracy measures based on classification probabilities for ups classification algorithm using leave-one-out cross-validation on the reference database.**

| | OA | kappa | RMSE |
|---|---|---|---|
| Existing pipeline | 0.582 | 0.335 | 0.375 |
| Our method | 0.743 | 0.576 | 0.377 |

In summary, we see that our method provides good accuracy, with the ability to distinguish between upsB and upsC groups, and provides further information with classification probabilities.

**Data S2.2.3 Thresholding probabilities to improve accuracy**

We further explored the possibility of applying a threshold to the classification probabilities from our method, so that sequences with the highest classification probability below the threshold would be considered unclassified.

In Figure III below, we consider the sensitivity and precision for the three groups for a range of thresholds on a logarithmic scale from 1 (the maximum possible threshold). We see that the accuracy of upsA classification is always high, as expected; meanwhile, both upsB and upsC accuracy improves with a higher threshold up to a point, after which upsB accuracy declines. This suggests that we can get better accuracy by applying a threshold, with the optimal threshold around 1-exp(-8). At this threshold, we show a significant improvement on all three accuracy measures used in the previous section (OA 0.899, kappa 0.843, RMSE 0.259).

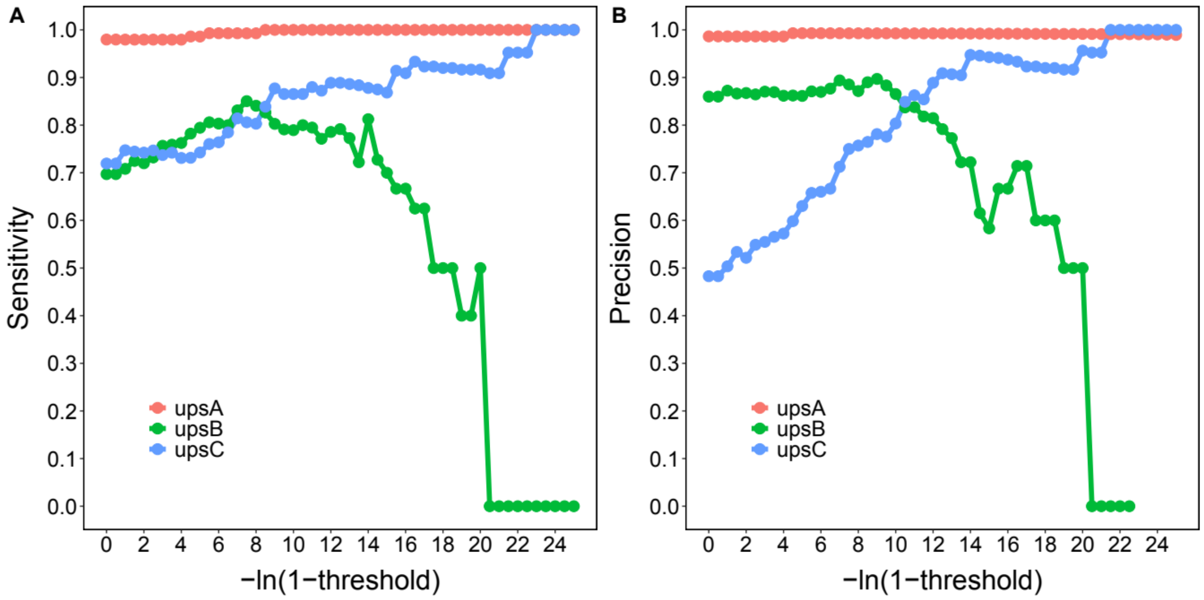


**Figure III. Sensitivity and precision of the ups classification algorithm for varying thresholds on the maximum classification probability.**

The drawback is that the number of classifications declines steeply with increasing threshold, as shown in Figure IV below. At the threshold of 1-exp(-8), we are only able to classify 22.5% of upsB and 33.7% of upsC sequences.

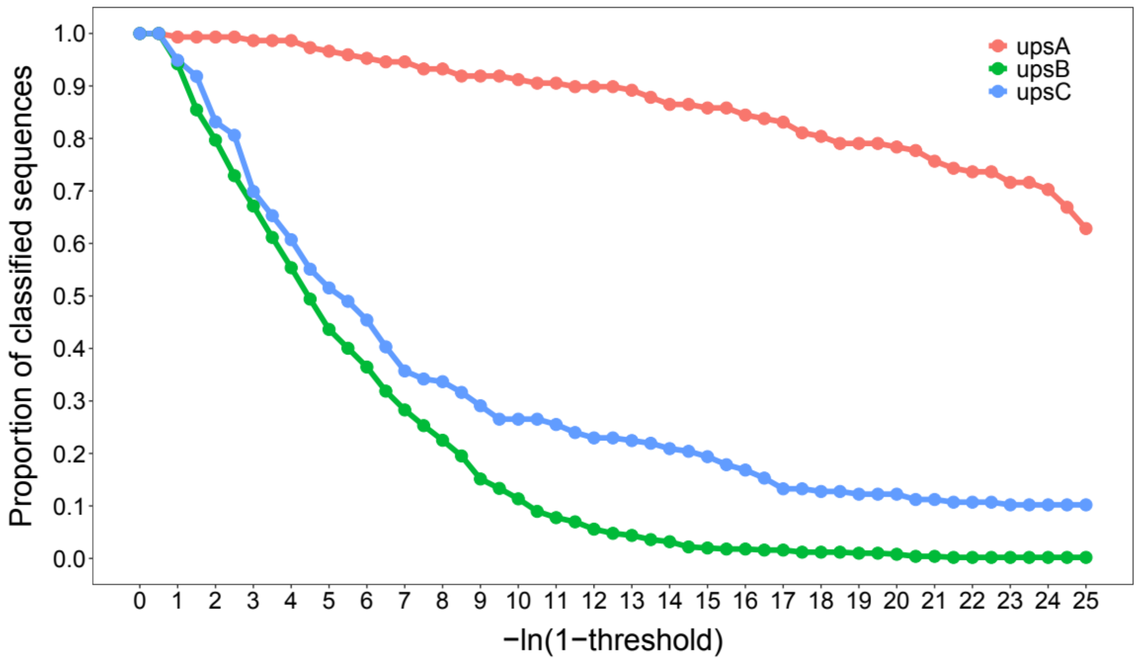


**Figure IV. Proportion of classified sequences for varying thresholds on the maximum classification probability.**

We conclude that thresholding is likely to be useful in certain cases, such as when studying a small number of particular types for which we want accurate classifications. However, we do not recommend its use when studying broad patterns in the data (as we do in this paper).
