## Supplementary material for "A paradoxical population structure of *var* DBLα types in Africa": Data S3

### Data S3. Study of DBLα conservation in Africa

**Table I. Dataset sources in Africa and Asia.** All datasets were used in the clustering step to generate representative DBLα types. However, further study on DBLα conservation was limited to only African locations with ≥ 100 isolates. For GhanaMRS, further study on DBLα conservation was limited to only surveys 1 to 7, excluding the pilot study.

| Continent | Location | Year(s) collected | DBLα source^ | # Isolates (initial) | # Isolates (≥20 types) | # Sites + | # Years + |
|---|---|---|---|---|---|---|---|
| Africa | Senegal (West)[1]* | 2001-2014 | Assembled *var* | 223 | 63 | | |
| Africa | Gambia (West)[1]* | 2008, 2013-2015 | Assembled *var* | 277 | 62 | | |
| Africa | Guinea (West)[1]* | 2011 | Assembled *var* | 197 | 94 | | |
| Africa | Mali (West)[1]* | 2007, 2012-2014 | Assembled *var* | 449 | 75 | | |
| Africa | GhanaMRS (West)[2] | 2012 (Pilot study) | AmpSeq | 161 | 158 | | |
| Africa | GhanaMRS (West)[2] | 2012-2017 | AmpSeq | 3,955 | 3,387 | 1 | 7 |
| Africa | Ghana (West)[1] | 2009-2015 | Assembled *var* | 1,041 | 475 | 2 | 5 |
| Africa | Nigeria (West)[1] | 2012 | Assembled *var* | 3 | 3 | | |
| Africa | Gabon (West)[3] | 2000 | AmpSeq | 200 | 176 | 1 | 1 |
| Africa | Democratic Republic of Congo (Central)[1]* | 2012-2015 | Assembled *var* | 366 | 92 | | |
| Africa | Malawi (Central)[1] | 2011 | Assembled *var* | 371 | 246 | 2 | 1 |
| Africa | Uganda (East)[4] | 2006-2007 | AmpSeq | 517 | 499 | 6 | 2 |
| Africa | Kenya (East)[1]* | 2007, 2009, 2014 | Assembled *var* | 131 | 26 | | |
| Asia | Bangladesh[1]* | 2012 | Assembled *var* | 49 | 46 | | |
| Asia | Cambodia[1]* | 2008-2012 | Assembled *var* | 623 | 439 | | |
| Asia | Laos[1]* | 2011-2012 | Assembled *var* | 82 | 79 | | |
| Asia | Myanmar[1]* | 2011-2013 | Assembled *var* | 58 | 56 | | |
| Asia | Thailand[1]* | 2011-2013 | Assembled *var* | 142 | 135 | | |
| Asia | Vietnam[1]* | 2009-2011 | Assembled *var* | 151 | 115 | | |

[1] (1), [2] (2), [3] (3), [4] (4)

^ "AmpSeq" refers to amplicon sequencing of the DBLα tag region; "Assembled *var*" refers to DBLα tags extracted from assembled *var* genes.

* Locations were excluded from the study either due to location (Asia is excluded) or due to limitations in the dataset size (i.e., in Africa but # Isolates (final) < 100).

\+ Values shown only for datasets retained in final spatial analysis, following the exclusion of individual isolates with < 20 DBLα types and subsequent exclusion of locations with < 100 isolates. "# Sites" and "# Years" refers to specific area and time sampling was performed, based on metadata available by MalariaGen e.g., Pf3k, Pf6 (5).

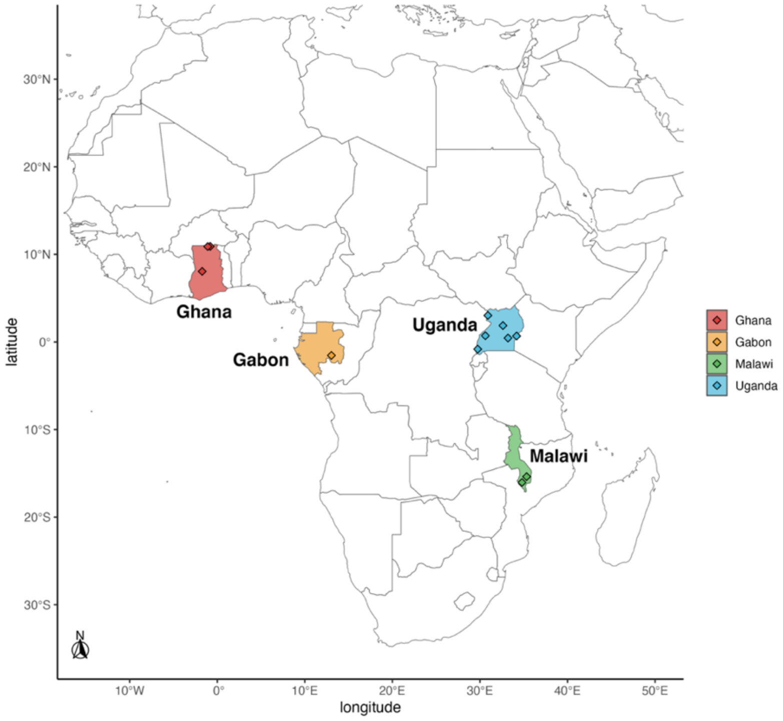


**Figure I. Final locations (i.e., countries) included in the spatial analysis of DBLα conservation in Africa.**

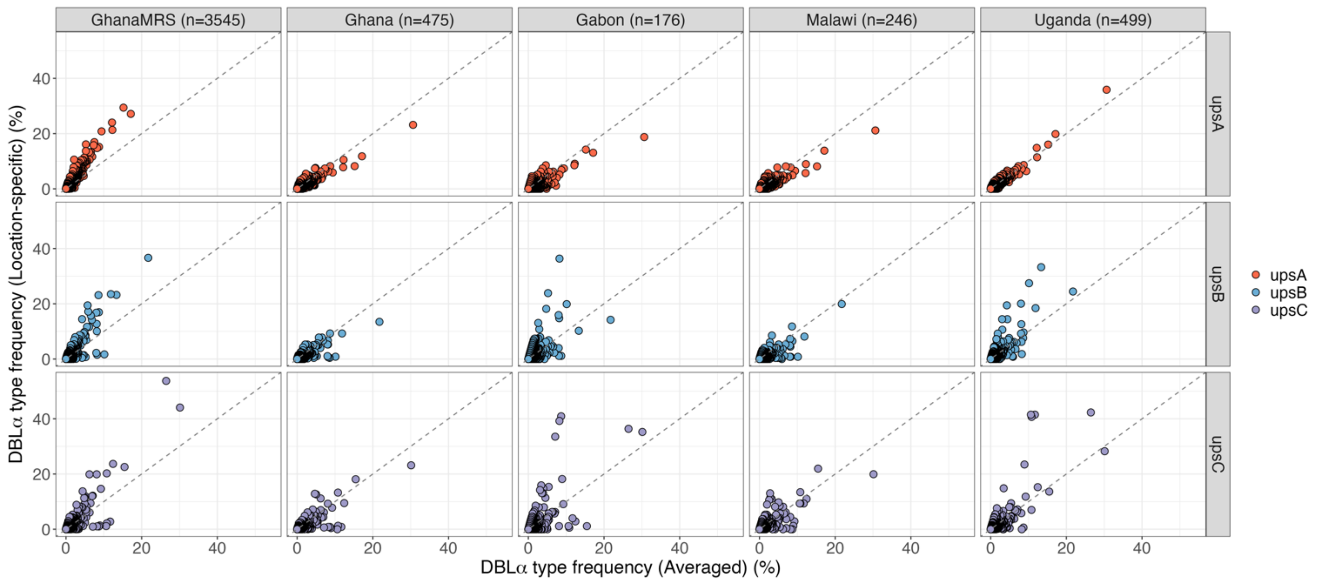


**Figure II. Conservation of DBLα types and frequencies at local and continent levels in four locations in Africa [Africa spatial analysis].** Location-specific frequencies (y-axis) are plot against location-averaged frequencies (x-axis), showing positive correlation between both frequencies. Points represent individual DBLα types, coloured by ups groups.

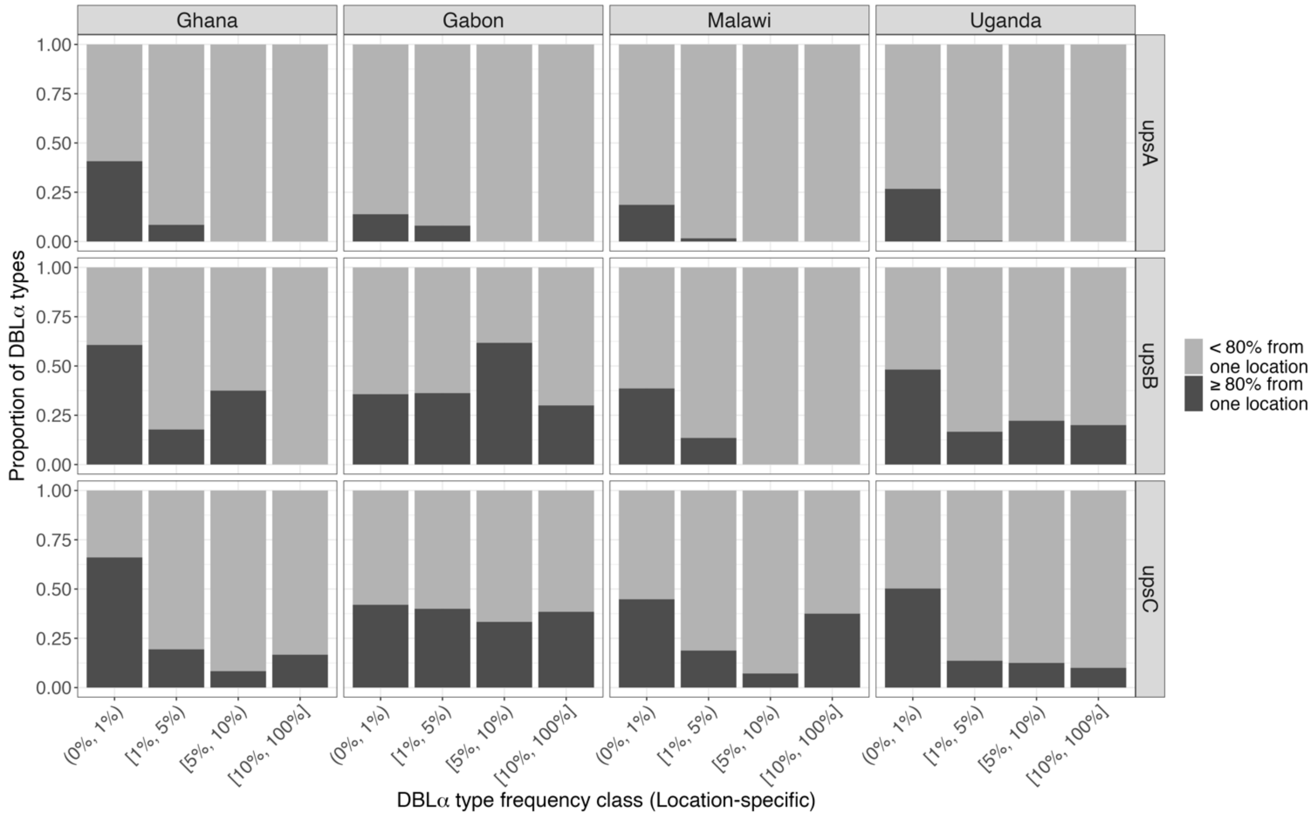


**Figure III. Proportion of DBLα types where a single location is a major source contributor (i.e. ≥80% of individuals for the DBLα type is from one location) [Africa spatial analysis].** The Ghana dataset includes DBLα types present in the combined datasets of "GhanaMRS" and "Ghana".

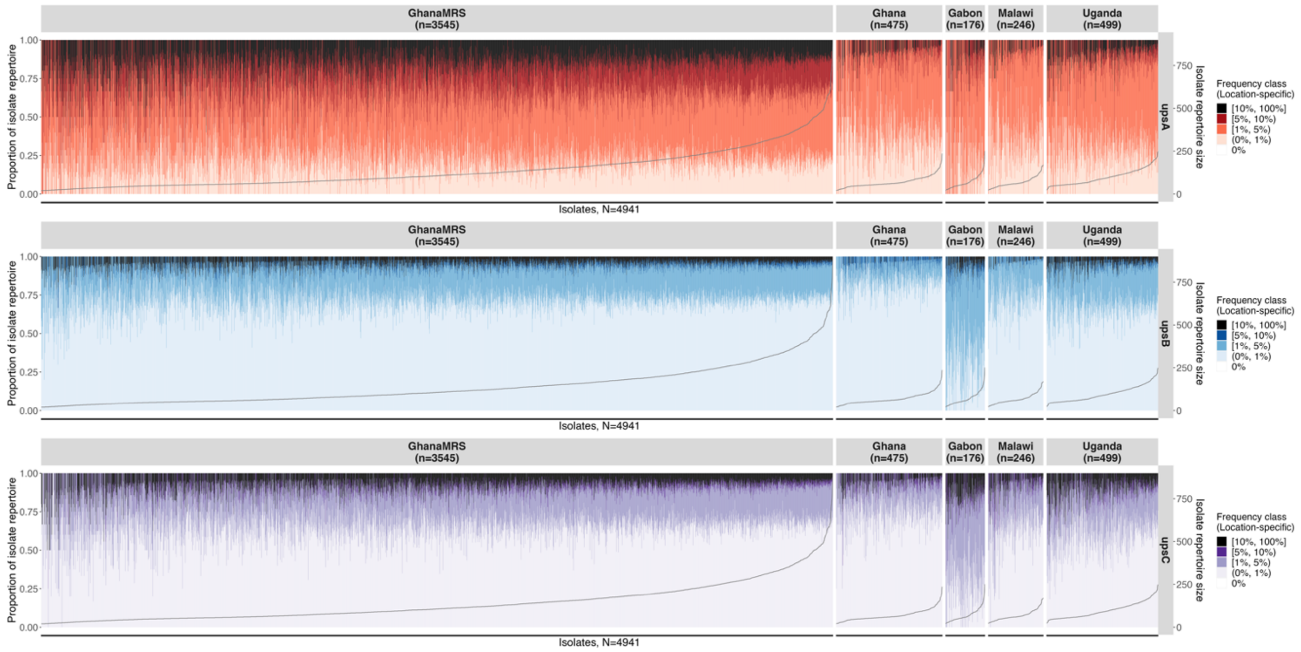


**Figure IV. Per-isolate frequency profiles show the composition of location-specific frequency classes in every isolate repertoire [Africa spatial analysis], using DBLα tag sequences extracted from assembled *var* gene data**. Vertical bars represent individual isolates, showing the composition of frequency classes in an isolate (left y-axis) by ups group (horizontal panels). Isolates are sorted by isolate repertoire size in increasing order, with isolate repertoire sizes indicated by the grey line (right y-axis).

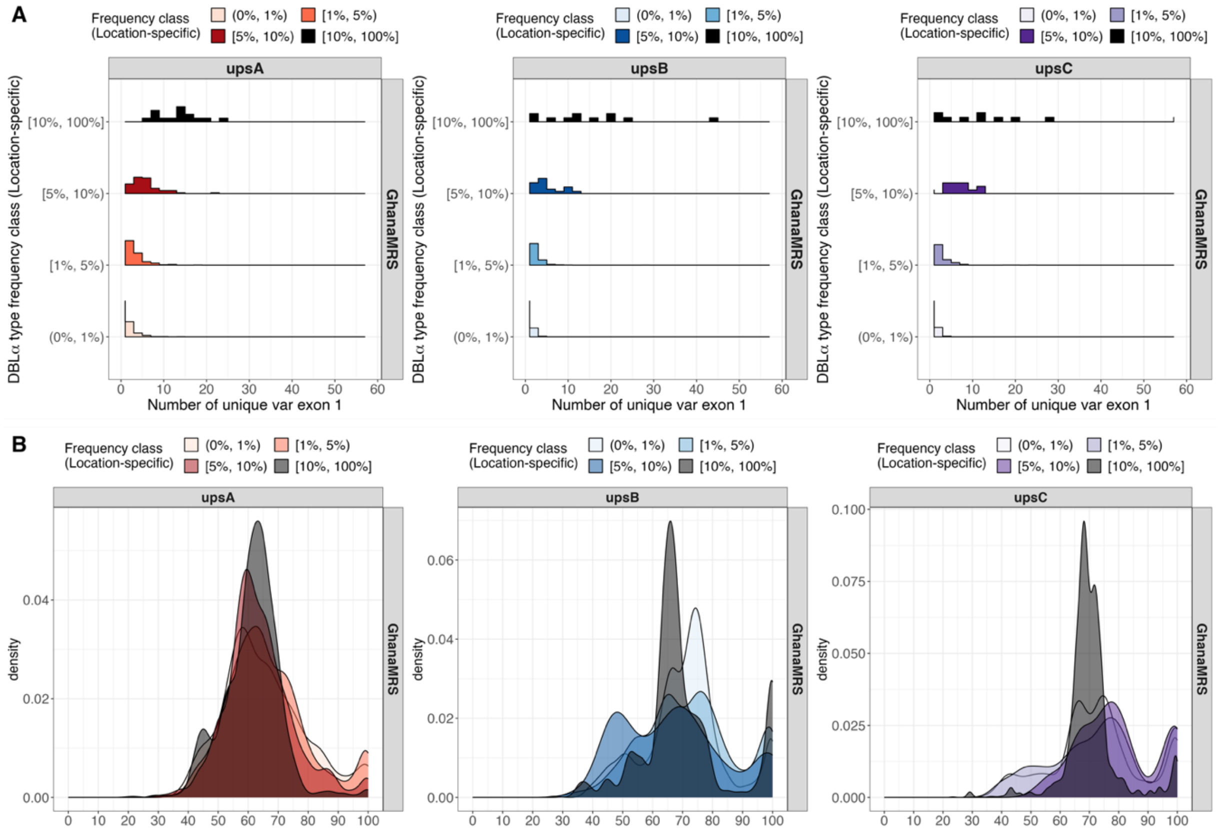


**Figure V. Relationship between DBLα types and *var* exon 1 for DBLα types found in the GhanaMRS data [Africa spatial analysis].** (A) Distributions show the number of unique *var* exon 1 sequences (in Navrongo) containing the same DBLα type (B) Distribution of nucleotide identities of pairs of aligned *var* exon 1. Horizontal panels represent the different location-specific frequency classes.
